# Eco-evolutionary theory and insect outbreaks

**DOI:** 10.1101/088047

## Abstract

Eco-evolutionary theory argues that population cycles in consumer-resource interactions are partly driven by natural selection, such that changes in densities and changes in trait values are mutually reinforcing. Evidence that this theory explains cycles in nature, however, is almost nonexistent. Experimental tests of model predictions are almost always impossible because of the long time scales over which cycles occur, but for most organisms, even tests of model assumptions are logistically impractical. For insect baculoviruses in contrast, tests of model assumptions are straightforward, and baculoviruses often drive outbreaks of forest-defoliating insects, as in the gypsy moth that we study here. We therefore used field experiments with the gypsy moth baculovirus to test two key assumptions of eco-evolutionary models of host-pathogen population cycles, that reduced host infection risk is heritable and costly. Our experiments confirm the two assumptions, and inserting parameters estimated from our data into the models gives cycles closely resembling gypsy moth outbreak cycles in North America, whereas standard models predict unrealistic stable equilibria. Our work shows that eco-evolutionary models are useful for explaining outbreaks of forest insect defoliators, while widespread observations of intense selection imposed by natural enemies on defoliators, and frequent laboratory observations of heritable and costly resistance in defoliators, suggest that eco-evolutionary dynamics may play a general role in defoliator outbreaks.

## Introduction

Eco-evolutionary theory has shown that natural selection can help drive cycles in predator-prey and other consumer-resource interactions, such that changes in trait values lead to changes in population densities, and vice versa (Abrams 2000). Recent work has focused in particular on the case for which selection by the consumer drives selection in the resource (Ellner 2013), so that increased consumer attacks lead to both decreases in resource densities and increased resource resistance, due to natural selection for increased resistance, while decreased attacks lead to both increased resource densities and reduced resistance, due to a fitness tradeoff between resistance and fecundity. Changes in population densities and changes in trait values are thus mutually reinforcing.

Eco-evolutionary cycles have been observed in models of predators and prey (Abrams and Matsuda 1997; Doebeli 1997; Ellner et al. 2011; Schreiber et al. 2011), hosts and parasitoids (Sasaki and Godfray 1999), and hosts and pathogens (Dieckmann 2002), but whether the models can explain population cycles in nature is unclear. Microcosm experiments have shown that ecoevolutionary predator-prey cycles occur in the laboratory (Fussmann et al. 2000; Yoshida et al. 2003), but laboratory conditions are often very different from field conditions, and field tests of the theory are effectively nonexistent (Abrams 2000). Here we therefore test eco-evolutionary theory using field data for the gypsy moth *(Lymantria dispar)* and its baculovirus.

Cycles of the gypsy moth and other outbreaking insects occur over time scales of decades and spatial scales of thousands of square kilometers, making full-scale experimental tests of model predictions impractical (Liebhold and Kamata 2000). We therefore first tested model assumptions, specifically the assumptions that resistance is heritable and costly. Notably, resistance in the models is defined in terms of overall infection risk (Elderd et al. 2008), whereas evidence of heritable and costly resistance in baculoviruses comes mostly from laboratory experiments that measure only infection risk given exposure (Boots and Begon 1993; Cory and Myers 2009; Watanabe 1987). Experiments with other host-pathogen interactions have similarly considered only infection risk given exposure (Altizer et al. 2003). Previous work has therefore not provided robust tests of model assumptions.

An underlying problem is that overall infection risk is best measured in the field, but for most host-pathogen interactions, field experiments are impractical. Meanwhile, for the few host-pathogen interactions for which experiments have measured infection risk in the laboratory or the greenhouse (Auld et al. 2014, 2013; Henter and Via 1995; Herzog et al. 2007; Zbinden et al. 2008), there are no data demonstrating that population cycles occur in nature. For insect baculoviruses in contrast, field experiments are straightforward (Elderd 2013), and because of the economic importance of the gypsy moth as an outbreaking forest pest, there are extensive data documenting gypsy moth population cycles (Johnson et al. 2005).

Previous efforts to explain these cycles, however, have met with limited success. Classical insect-pathogen models require variability in host infection risk to prevent pathogen extinction, but realistically high variability causes the models to produce a stable equilibrium instead of cycles (Dwyer et al. 2000). Extending classical models to allow pathogen transmission to be affected by induced plant defenses leads to models that can explain gypsy moth cycles, but the resulting host-pathogen/induced-defense models require particular spatial configurations of tree species (Elderd et al. 2013), and so cannot explain outbreaks in some forest types in which gypsy moth outbreaks have been observed to occur (Haynes et al. 2009).

Allowing for heritable variation in insect-outbreak models can also lead to realistic cycles when variation is high, irrespective of forest type (Elderd et al. 2008), suggesting that ecoevolutionary models may provide a better explanation for gypsy moth outbreak cycles. For mathematical convenience, however, the models in question made the unrealistic assumption that heritability is perfect, even though heritability is undoubtedly less than one. Moreover, reduced heritability is strongly stabilizing in predator-prey models (Schreiber et al. 2011), suggesting that low heritability may prevent cycles in insect-outbreak models. Here we therefore test whether eco-evolutionary dynamics can explain gypsy moth outbreaks, first by extending previous models to allow for imperfect heritability, second by estimating the heritability and costs of increased resistance, and third by testing whether our heritability and cost estimates produce model cycles that match cycles seen in nature.

Eco-evolutionary microcosm models show 1/2-cycle lags between predator and prey population peaks that qualitatively differ from the 1/4-cycle lags of classical models (Fussmann et al. 2000; Yoshida et al. 2003), whereas population cycles in our model closely resemble the cycles of classical, non-evolutionary models (Dwyer et al. 2000). This difference between our models and microcosm models apparently occurs because our models include discrete generations, a basic feature of the biology of many outbreaking insects (Hunter 1991), emphasizing the importance of constructing models from field data. The similarity of our model’s predictions to those of classical models, however, means that, unlike with microcosm models, it is not possible to test model predictions through qualitative comparisons to data.

We therefore instead tested model predictions by comparing the period and amplitude of the models (Kendall et al. 1999) to the period and amplitude seen in data for gypsy moths (Johnson et al. 2005). Because the predictions of our model match the data, our work supports the hypothesis that eco-evolutionary consumer-resource cycles occur in nature, not just in microcosms. Moreover, because the gypsy moth is a major pest of hardwood forests in North America (Elkinton and Liebhold 1990), and because the virus plays a role in gypsy moth control (Podgwaite et al. 1993), our model may be useful for guiding gypsy moth management, as we discuss.

## Methods

### Baculovirus Biology and Eco-Evolutionary Models of Insect Outbreaks

In forest-defoliating insects like the gypsy moth, baculovirus transmission occurs when larvae accidentally consume infectious particles or “occlusion bodies” during feeding, and so only larvae can become infected (Cory and Hoover 2006). Larvae that consume enough occlusion bodies die, after which viral enzymes break down the larval cuticle, releasing occlusion bodies into the environment (Elderd 2013). In high density populations, multiple rounds of transmission cause severe mortality, terminating outbreaks (Moreau and Lucarotti 2007). The virus then overwinters by contaminating the egg masses produced by surviving insects (Murray and Elkinton 1989).

This simple biology can be accurately described by a Susceptible-Exposed-Infected-Recovered (SEIR) model, modified to allow for host variation in infection risk (Dwyer et al. 2002,1997):

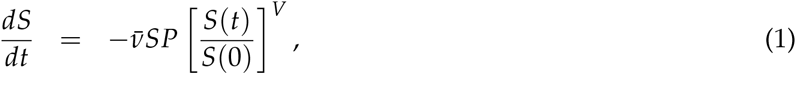

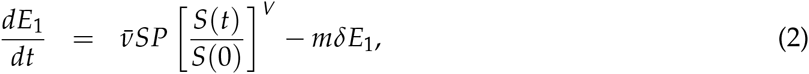

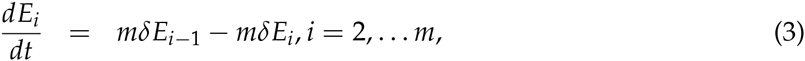

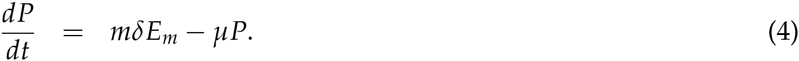

Here, *S* and *P* are the densities of healthy hosts and infectious cadavers, respectively, while *E_i_* is the density of exposed but not yet infectious hosts, so that *i* indicates the exposure class. Allowing for *m* exposure classes produces a distribution of times to death, with mean 1/*δ* and coefficient of variation (C.V.) 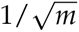 (Keeling and Rohani 2008). Variation in infection risk is described by the transmission term 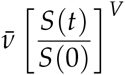, such that the initial mean transmission rate is and the squared C.V. of transmission rates is *V* (Dwyer et al. 2000).

Transmission 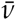 determines host infection risk, and so including variation in transmission is a first step in allowing natural selection to drive the evolution of host infection risk. The second step is to allow for multiple host generations by including host reproduction, which we accomplish by embedding equations (1)-(4) in a discrete-generation model (see Appendix A). Because generalist predators and parasitoids also affect gypsy moth populations, we include a term that describes the fraction surviving predation (Dwyer et al. 2004):

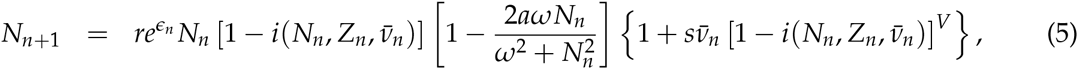

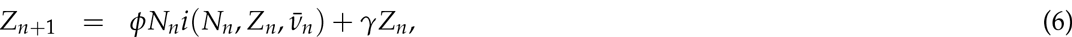

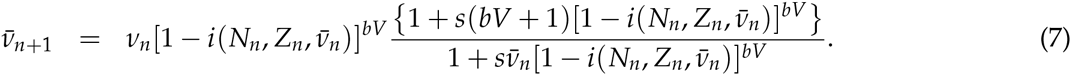

The resulting model tracks generational change not just in hosts *N_n_* and pathogens *Z_n_*, but also in the average transmission rate 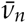, which is the infection risk at the beginning of generation *n* (for simplicity, we assume constant *V*). The fraction infected during the epizootic 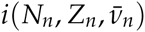 is calculated using equations (1)-(4), assuming an eight week epizootic (Fuller et al. 2012).

Host density *N*_*n*+1_ in host generation *n* + 1 is the product of baseline fecundity *r*, a stochasticity term *e^εn^*, host density in the preceding generation *N_n_*, and the fraction surviving the epizootic 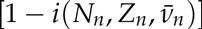. The stochasticity parameter *ε_n_* is a normal random variate with mean 0 and standard deviation σ, which has a different value in each generation, representing the effects of weather (Williams et al. 1990). Generalist predation is represented by the term 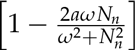 which describes host survival as determined by a Type III functional response (Dwyer et al. 2004). At low host *N_n_* and pathogen *Z_n_* densities, reproduction increases linearly with increasing infection risk 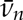, according to 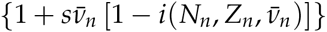, reflecting a fecundity benefit of increased risk that is reduced when the infection rate 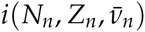 is high due to high *N_n_* and/or high *Z_n_*. Changes in host density are thus partly driven by balancing selection, such that higher infection risk leads to increased mortality but also increased fecundity.

Pathogen density *Z_n_+1* in host generation *n* + 1 is equal to the density of infectious cadavers produced in the preceding generation’s epizootic, 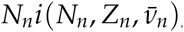, times the effective overwintering rate *ϕ*. Because *ϕ* allows for the higher susceptibility of neonate larvae, which are orders of magnitude more susceptible than later stage larvae, previous work has shown that *ϕ* > 1 (Fleming-Davies and Dwyer 2015; Fuller et al. 2012). Long-term pathogen survival is represented by the cadaver density *Z_n_* times the long-term pathogen survival rate *γ* (Fuller et al. 2012).

Infection risk 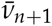 in generation *n* + 1 is equal to the preceding generation’s risk 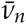 times the fraction infected 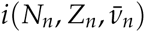, reflecting selection for reduced risk. Risk also increases because of the cost of resistance, due to linear increases in fecundity with increases in previous-generation risk when host and pathogen densities are low, 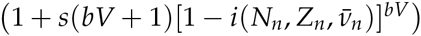, an effect that is again reduced by high infection rates. As with host density, infection risk is thus determined by balancing selection, except that risk depends only on the scaling parameter *s* and not the baseline fecundity *r*. Finally, *b* is the heritability of risk, so that *bV* is the fraction of variation that is due to additive genetic factors. High values of heritability *b* thus strengthen the effects of selection.

Population cycles in this model occur because of the consumer-resource interaction between the host and the pathogen, and because of natural selection on infection risk and fecundity (fig. 1). Low virus mortality allows high-fecundity, high-infection-risk genotypes to rise in frequency, leading to increasing virus density and increasing average infection risk, which in turn cause virus epizootics that decimate the host population. Host density is then low for multiple host generations not just because of the pathogen and the generalist predators, but also because the survivors of virus epizootics are more resistant to the pathogen and therefore suffer a fecundity cost. Eventually, however, falling virus density and rising fecundity together drive increases in host density, leading to a new outbreak. Natural selection therefore combines with ecological factors to drive outbreaks. Moreover, models in which there is no generalist predator, or in which the pathogen is instead a parasitoid, give qualitatively similar results (see Online Appendix).

**Figure 1:**
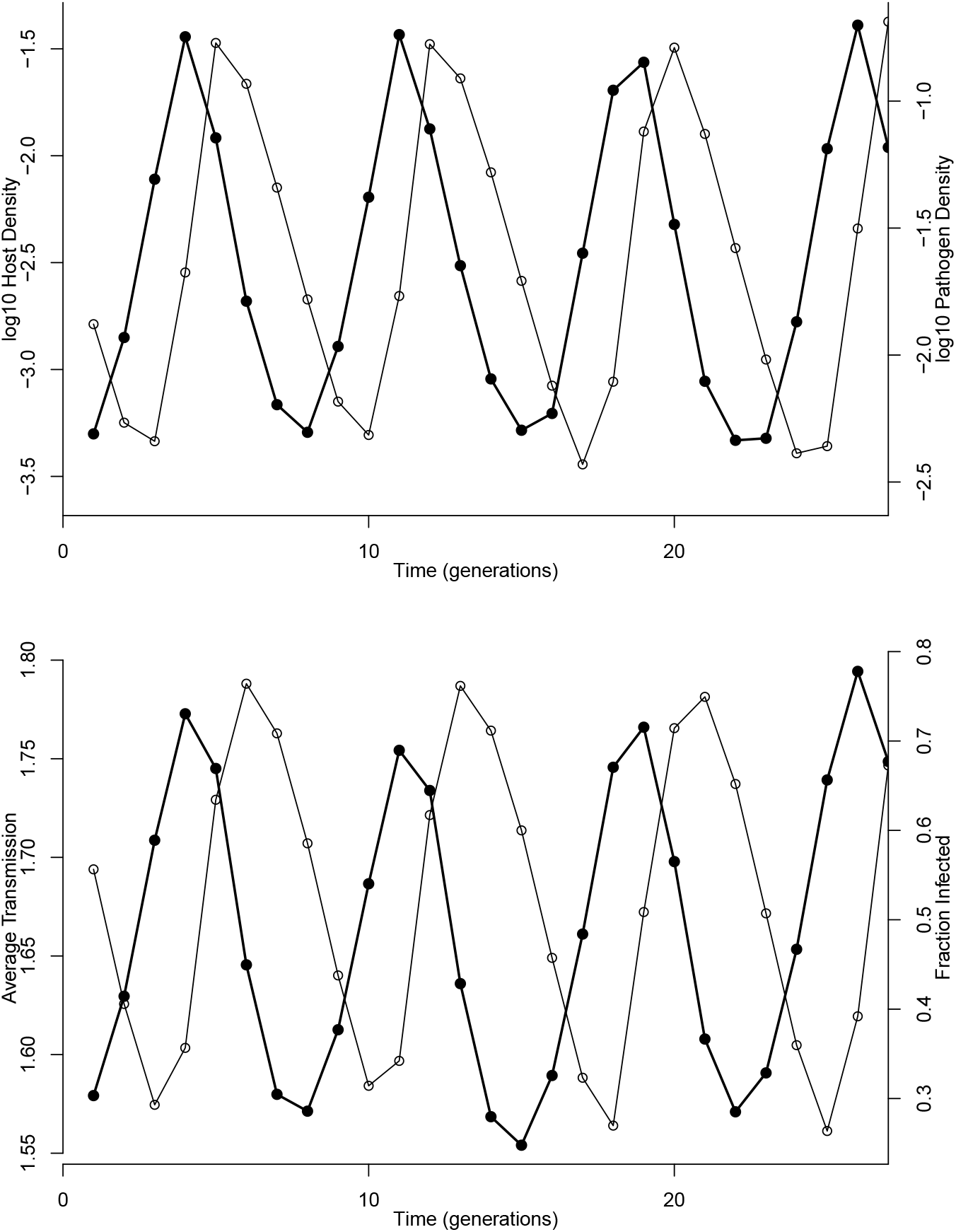
A single realization of the model equations (5)-(7). The top panel shows changes in host densities *Z_n_* and pathogen densities *Z_n_* (black points/bold lines and open points/gray lines, respectively), while the bottom panel shows the corresponding changes in average infection risk 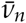 and fraction infected *i* (black points/bold lines and open points/gray lines, respectively). Heritability *b* = 0.126, fecundity cost on density *r* = 0.21, cost scaling parameter *s* = 1.21, total variation *V* = 2.97, the median parameter values calculated from our experimental data. Pathogen overwintering parameter *ϕ* = 7.4 and long-term survival *γ* = 0.3 from Fuller et al. (2012) (*γ* is on the high end of reasonable values, but variation in *γ* has only modest effects, see Supporting Information), generalist predation parameters *a* = 0.96, *ω* = 0.14 from Dwyer et al. (2004).

### Field Experiments to Estimate the Heritability and Cost of Reduced Infection Risk

The key assumptions of our eco-evolutionary model are that infection risk *ν* is heritable, so that heritability *b* > 0, and that there is a cost of reduced risk, so that the relationship between fecundity and risk has positive slope *rs* > 0. Previous work produced preliminary evidence that gypsy moth infection risk is heritable, without estimating heritability, and without providing evidence that reduced risk is costly (Elderd et al. 2008). Our experiments were therefore designed both to test whether infection risk is heritable and costly, and to estimate the heritability and cost parameters, to determine if the parameters fall in the right range to produce realistic outbreaks in the model.

Part of the reason why it is important to measure infection risk in the field is because infection risk depends on feeding behavior, which cannot be easily allowed for in laboratory experiments (Dwyer et al. 2005). Feeding behavior affects the overall risk of infection by determining the risk of exposure, with the proviso that the overall risk of infection is also affected by the risk of infection given exposure, which is instead determined by the insect’s immune response (Páez et al. 2015). Feeding behavior is thus important because gypsy moth larvae can sometimes detect and avoid virus-infected cadavers (Capinera et al. 1976), a heritable behavior (Parker et al. 2010), but variation in risk given exposure is also heritable (Páez et al. 2015). By measuring overall infection risk, we thus allowed for variation in both components of infection risk.

To prevent larval emigration and virus decay (Fuller et al. 2012), all foliage was covered in mesh bags. Also, the exposure period was short enough that no new virus deaths occurred during either experiment, and so there was no change in the density of virus after the experiment began. By setting the change in virus density 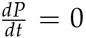, we can then simplify equations (1)-(4) to produce an expression for the fraction infected *i* at the end of the experiment (Dwyer et al. 2000):

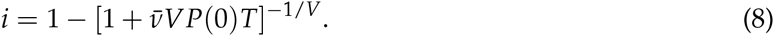

Here *P*(0) is the density of virus-infected cadavers, and *T* is the length of time for which the experiment runs. Because in our experiments we measured *i*, it was possible to use nonlinear fitting routines (see Online Appendix) to fit equation (8) to our data, and thus to estimate average infection risk 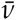 and variation *V*. In previous work, inserting such experimental parameter estimates into equations (1)-(4) produced infection rates that are close to those seen in nature (Dwyer et al. 2002,1997; Dwyer and Elkinton 1993), suggesting that our protocol produces accurate parameter estimates.

Inferences about transmission parameters are stronger in experiments that include a range of virus densities, and so we used densities of 0, 25, 50, and 75 virus-infected cadavers per 40-leaf branch (Elderd et al. 2008). After larvae had fed on virus-contaminated foliage in the field, we reared them in individual diet cups in the lab until death or pupation. Infected larvae are usually easily recognizable because the virus causes larvae to disintegrate, but in cases of uncertainty, we examined smears from dead larvae for the presence of occlusion bodies, which are easily visible at 400 x (Fleming-Davies et al. 2015). Because we used an area in which gypsy moth densities were very low, and because all eggs were surface-sterilized in dilute formalin (Dwyer and Elkinton 1995), infection rates on uninfected control foliage were low (heritability experiment: 12/209 = 5.7%; cost experiment: 6/371 = 1.6%), and so we do not consider controls further.

To estimate heritability, we decomposed the variance in infection risk 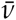 into additive genetic variance and environmental variance. Additive genetic variance can be estimated from the variance in 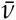 across half-sibling groups, which is due to sire effects *S_i_*. Maternal effects in the gypsy moth can arise from variability in egg provisioning (Diss et al. 1996), but there may also have been small-scale differences between rearing cups. We therefore collectively denote maternal and small-scale rearing effects as *M_j_*. Also, for logistic reasons, larvae were not all deployed in the field on the same day, and so we also allowed for environmental variance due to a start-day effect *D_k_*. We thus expressed average infection risk in terms of sire, dam and day effects, so that 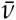 in equation (8) becomes:

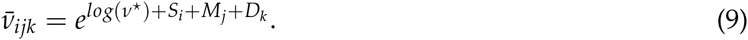

Here *v*\* is the baseline infection risk. Following quantitative genetic theory (Falconer and Mackay 1996), heritability is then estimated as;

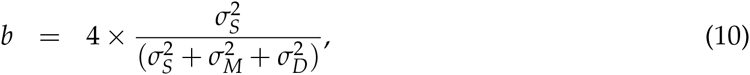

such that 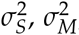, and 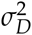 are the variances due to the sire, maternal and day effects, respectively, which we estimated using a hierarchical Bayesian fitting routine. To test whether variation in infection risk is heritable, we then used AIC analysis to choose between models that did or did not include sire effects, maternal effects and day effects.

In principle, variation in infection risk could be explained solely by sire, dam and day effects, which would mean that we could allow *V* → 0 in equation (8) (Dwyer et al. 2005). In addition, however, there is small-scale spatial variation in virus density (Eakin et al. 2015), which, in contrast to sire, dam and day effects, does *not* depend on factors that vary between host families. Because of this variation, allowing *V* → 0 in equation (8) provided a much worse fit to the data, and so we instead assumed *V* > 0.

To estimate the cost of reduced infection risk, we again fit equation (8) to data for each full-sibling group, and we regressed average female pupal weight in each group on the average infection risk in that group (Elderd et al. 2008). Because pupal weight is strongly correlated with egg number (Páez et al. 2015), a positive relationship between pupal weight and infection risk indicates that there is a fecundity cost of reduced risk.

## Results

In our heritability experiment, infection rates and infection risk varied strongly across half-sibling families (fig. 2A, B), with 13% of the variation in risk explained by additive genetic variation (*b* = 0.13). AIC analysis then showed that models with sire effects explain the data vastly better than models without sire effects (ΔAIC = 295.6 for the best model without a sire effect, Table 1). The 95% highest posterior density interval (Bayesian equivalent of a 95% confidence interval) on heritability *b* was broad (HPD = 0.0013,0.48), but excluded values below 10^-3^. We therefore conclude that infection risk in the gypsy moth has low but non-zero heritability, confirming the first key assumption of our eco-evolutionary model.

**Table 1:** AIC analysis of transmission models. “MLL” is the maximum log likelihood. The best model, which allows for sire, dam, and day effects, is in bold face. AIC weights for all but the best model were all less than 10^-5^, and are therefore omitted.

| Model | Parameters | # parameters | MLL | $\Delta\text{AIC}$ |
| --- | --- | --- | --- | --- |
| **1** | $\nu, V, \text{Sire}_i, \text{Dam}_j, \text{Day}_k$ | **131** | **-1853.6** | **0.0** |
| 2 | $\nu, V, \text{Dam}_j, \text{Day}_k$ | 93 | -2038.4 | 295.6 |
| 3 | $\nu, V, \text{Sire}_i, \text{Dam}_j$ | 124 | -1866.3 | 11.4 |
| 4 | $\nu, V, \text{Sire}_i$ | 40 | -2073.8 | 258.4 |
| 5 | $\nu, V$, no random effects | 2 | -2100.0 | 234.8 |

**Figure 2:**
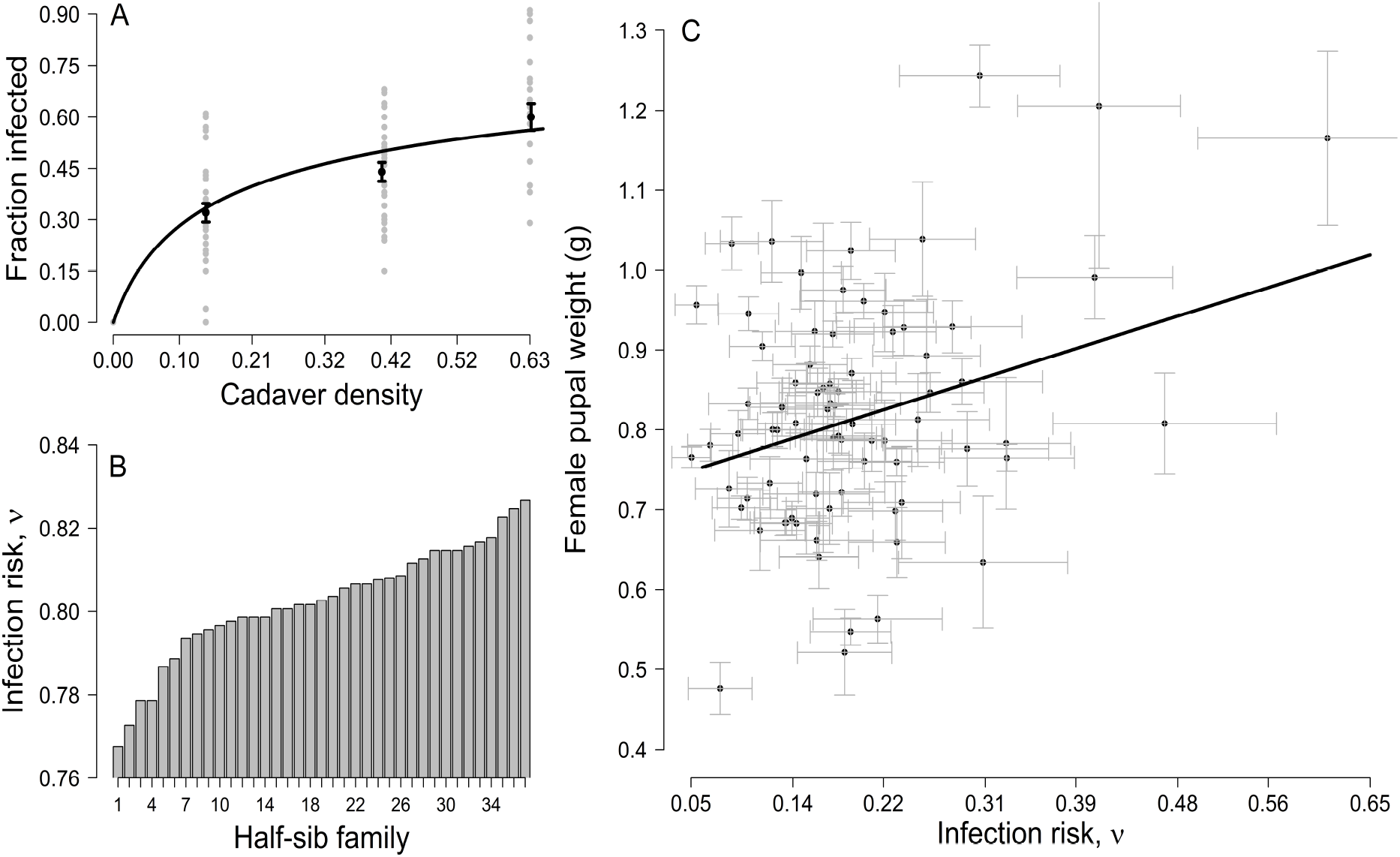
A) Relationship between fraction infected and density of virus-infected cadavers in our heritability experiment. Gray dots show the infection rate for each half-sibling family, demonstrating that there is meaningful variation across half-siblings, in turn suggesting that infection risk is heritable. The fitted curve is the best-fit version of the best transmission model, which includes sire, dam and day effects (Table 1). Median values for 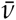 and *V* are 0.80 (HPD = 0.43, 1.37) and 2.97 (HPD =1.55, 4.59), respectively, which are close to values from previous experiments (Elderd et al. 2008). B) Variation in average infection risk 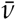 across half-sibling families, again suggesting that infection risk is heritable. C) Fecundity cost of resistance, as demonstrated by a positive relationship between infection risk and female pupal mass, a surrogate measure of fecundity (Páez et al. 2015).

In our cost experiment, there was a noisy but positive relationship between female pupal weight and infection risk (fig. 2C). The slope of the resulting regression line was significantly greater than zero, confirming that reduced infection risk has a fecundity cost (median intercept = 0.73, upper and lower 95th percentiles = 0.70, 0.80; median slope = 0.45, upper and lower 95th percentiles = 0.09,0.56, see Online Appendix for a description of how we bootstrapped the regression parameters to account for error in both infection risk and pupal weight.). In this regression, however, we included only survivors of virus exposure, which may have led to an underestimate of the cost. This is because variation between exposed individuals within groups could have led to an over-representation of low-risk/low-fecundity individuals among survivors.

We therefore carried out a second regression in which we also included insects that were reared in the lab but never exposed to the virus. The best model in this case assumed that the regression lines for exposed and unexposed individuals had the same slopes but different intercepts (see Online Appendix). The cost of reduced risk was thus indistinguishable for exposed and unexposed individuals, probably because variation within full-sibling groups was substantially smaller than variation between full-sibling groups. The difference in intercepts was likely due to differences in diet, because virus-exposed insects fed for a week on foliage, whereas lab-reared insects were fed only on artificial diet, which allows for higher growth (Rossiter 1991). In what follows, however, we use estimates of the cost parameters that are based only on survivors of the field experiment, because the pupal weights of field-reared insects were closer to pupal weights seen in nature (Páez et al. 2015). Irrespective of these complications, however, our data generally show that female gypsy moth larvae with reduced infection risk have reduced fecundity as adults, confirming the second key assumption of our eco-evolutionary model.

Because reduced infection risk is heritable and costly, balancing selection must inevitably play a role in determining infection risk in the gypsy moth, but that does not mean that selection will inevitably drive gypsy moth outbreaks. In particular, realistic outbreak cycles in our model do not occur for all parameter values. To test whether our parameters fall into the range that does give realistic cycles, we therefore inserted the parameters into the model to test whether the parameterized model can reproduce data on gypsy moth outbreak cycles.

In carrying out this test, it is important to note that cyclic population dynamics are usually at least moderately sensitive to initial densities, which are unknown. Kendall et al. (1999) therefore argue that a useful way to compare model cycles to data is by comparing periods and amplitudes, which in the long run are insensitive to initial conditions. For our estimates of the heritability and cost of infection risk, the average cycle period is 7.4 years, and the average amplitude is 2.1 orders of magnitude (fig. 1, note that we adjusted the cost parameters *r* and *s* to allow for non-disease, non-predation mortality, see Online Appendix).

Gypsy moth outbreaks in North America have had periods between 5 and 9 years (Johnson et al. 2005), and amplitudes between 2 and 4 orders of magnitude (Jones et al. 1998; Skaller 1985; Williams et al. 1990). Model periods and amplitudes thus fall within the range of values seen in nature, while variation around the median estimates has only modest effects (fig. 3). Furthermore, previous experiments showed that average infection risk falls in synchrony with falling population densities (Elderd et al. 2008), also as predicted by the model (fig. 1). The model predictions are thus confirmed by both observational and experimental data, suggesting that natural selection plays an important role in gypsy moth outbreaks.

**Figure 3:**
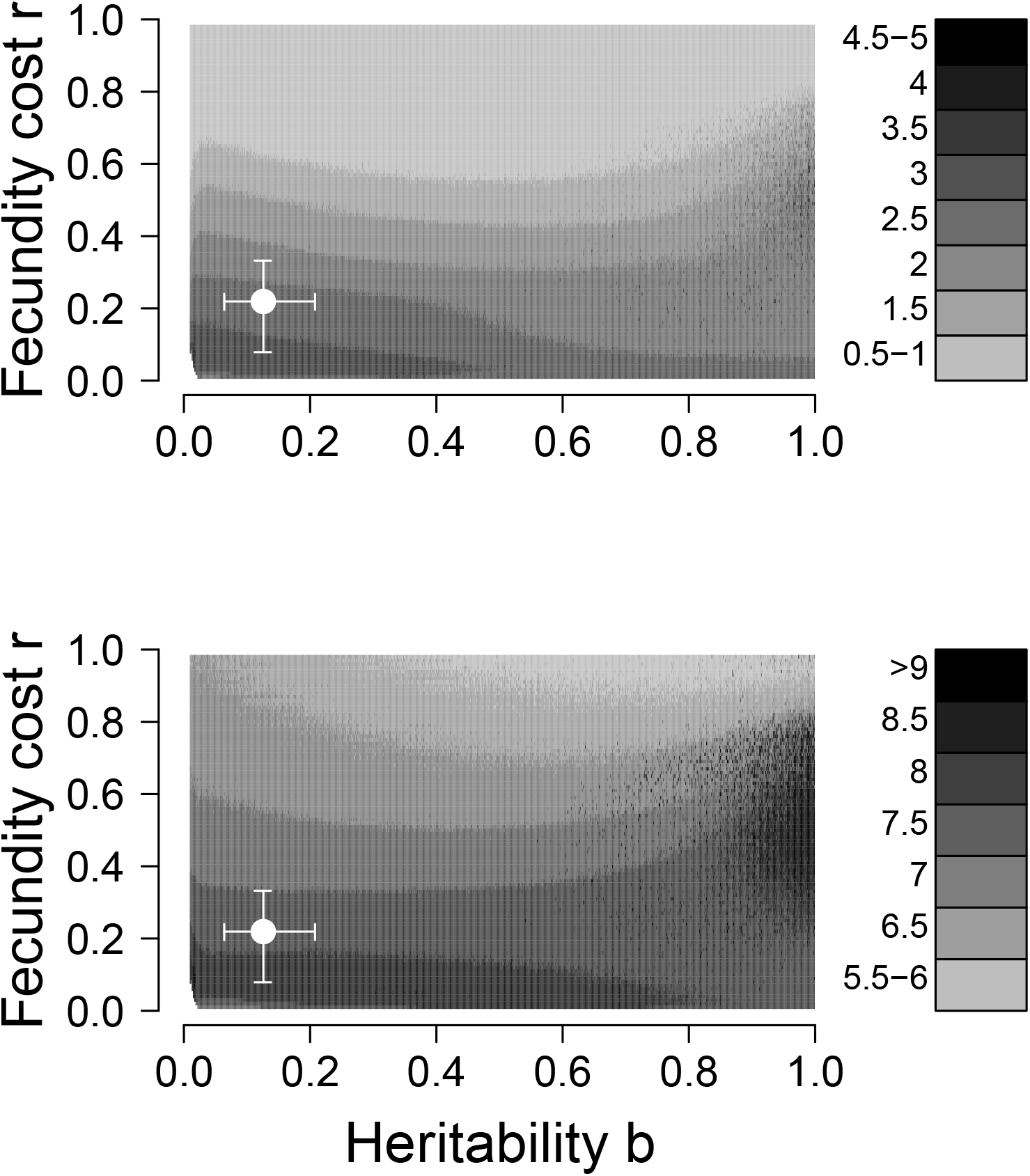
Effects of variation in baseline fecundity *r* and heritability *b* on the period and amplitude of outbreak cycles in the long term model, equations (5)-(7). Remaining parameters are as in Fig. 3. The top panel shows average cycle amplitudes in orders of magnitude, while the bottom panel shows the average period in years. The white dot represents our median estimates of *r* and *b*, with error bars indicating the interquartile range.

Given that our model provides a useful description of gypsy moth population cycles, we can use it to address an issue of general importance in eco-evolutionary theory, namely how heritability affects population stability. As fig. 3 shows, in our model intermediate heritability is at least mildly stabilizing, whereas in continuous-time eco-evolutionary models of predatorprey interactions, reduced heritability is instead stabilizing (Schreiber et al. 2011). In both types of models, high heritability causes such a rapid response to selection that infection risk undergoes large-amplitude fluctuations, driving severe fluctuations. In our model, however, low heritability is destabilizing because it leads to a slow response to selection, exacerbating the delayed density-dependence that drives population cycles. This difference likely occurs because our model includes the realistic assumption of seasonal reproduction, which allows for strong delayed density-dependence at low heritability.

A second general point is that our model apparently does not have either the 1/2-cycle lags (Yoshida et al. 2003) nor the cryptic population cycles (Yoshida et al. 2007) that often occur in microcosm models, instead showing the 1-2 generation lags that are apparent in fig. 1. We have not been able to prove that longer lags do not occur in our model, but for almost all of the parameter values in fig. 3, the average lag was two generations or less (99.6% of 96,981 parameter sets that did not cause host or pathogen extinction). This is important because the occurrence of 1-2 generation lags in our model is supported by data showing that baculovirus infection rates peak shortly after host densities in several outbreaking insects (Moreau and Lucarotti 2007; Moreau et al. 2005). The model prediction of lags shorter than 1/2-cycle is thus confirmed by the data.

Half-cycle lags in microcosms are more likely when costs are low, but reducing the cost-scaling parameter *s* in our model does not meaningfully change the lag (for *s* = 0.2, the lower bound on the HPD: 94.8% of 87,515 parameter sets; for *s* = 0.02: 91.3% of 13,387 parameter sets, note that reducing *s* tends to increase the probability of host and/or pathogen extinction, see Online Appendix). We therefore suspect that the reason why our models do not show half-cycle lags or cryptic cycles is again that our models include only one host generation per year, whereas in microcosms conditions are constant and breeding is therefore continuous (Hiltunen et al. 2014).

## Discussion

In arguing that natural selection helps drive gypsy moth population cycles, we note that Allstadt et al. (2013) showed that gypsy moth populations in North America cycled from 1943-1965, and from 1978-1996, but that there were periods of non-cyclic dynamics at other times (data points after 2009 were too few to permit testing, but more recent outbreaks suggest that cycling may have returned, G. Dwyer, pers. obs.). The same authors nevertheless showed that a temporary lack of cycles can be explained by a version of the Dwyer et al. (2004) model that includes stochastic fluctuations in generalist-predator density. Given that our model is essentially the Dwyer et al. model plus natural selection, we suspect that our model would show similar behavior. Allowing for fluctuating predator populations is nevertheless an important next step.

A related point is that Allstatdt et al., as well as other other researchers modeling gypsy moth outbreak data (Bjørnstad et al. 2010; Haynes et al. 2009; Walter et al. 2015), forced the nonevolutionary Dwyer et al. model to show cycles, by using estimates of host variation *V* that have been shown to be unrealistically low (Elderd et al. 2008). Low values of *V* prevent the unrealistic stable equilibria that occur when *V* is realistically high, but inferences based on incorrect models are often unreliable (Box 1979). A consideration of natural selection may thus be widely useful for understanding data on gypsy moth outbreaks.

More broadly, to focus on the effects of natural selection, we did not include three other mechanisms known to affect gypsy moth population dynamics in North America. First, the fungal pathogen *Entomophaga maimaiga*, introduced in 1990, often causes high mortality when rainfall is abundant (Hajek 1999), with the proviso that it may be only weakly density-dependent (Hajek et al. 2015; Liebhold et al. 2013). *E. maimaiga* therefore may have only mild effects on gypsy moth cycles, but its long term effects are not yet understood. Including both *E. maimaiga* and the baculovirus is therefore an important next step in modeling gypsy moth outbreak cycles.

Second, we assumed a monomorphic virus population, but in fact the gypsy moth virus is at least moderately polymorphic (Fleming-Davies et al. 2015). This is important because for simplicity we assumed that pathogen variation was constant, but including pathogen variability, and thus host-pathogen coevolution, could allow for the maintenance of host variation (Sasaki and Godfray 1999). Adding pathogen variation to our model is therefore a second important next step.

Lastly, as we mentioned earlier, the stabilizing effect of realistically high host variation, which in our models is mitigated by natural selection on infection risk, can also be mitigated by defoliation-induced defensive compounds, such that host-pathogen/induced-defense models can explain variability in outbreak periods between forest types that have intermediate versus high frequencies of oaks (Elderd et al. 2013). Host-pathogen/induced-defense models, however, do not allow for outbreaks in forest types with low frequencies of oaks, even though outbreaks have been observed in such forests (Haynes et al. 2009). Moreover, the models are sensitive to changes in the clumping of oaks within the forest (Elderd et al. 2013), whereas our eco-evolutionary models are insensitive to a range of changes in model structure (see Online Appendix). Both models nevertheless have extensive empirical support, and so including induced defenses is a third important next step. That said, our experimental results provide conclusive evidence that natural selection alters infection risk in the gypsy moth, and it is therefore likely that selection has at least some effect on outbreaks, confirming eco-evolutionary theory in a broader sense.

This list of other factors that may also affect gypsy moth cycles raises the larger point that, in the absence of outbreak-scale experiments, our work cannot provide conclusive proof for the importance of eco-evolutionary dynamics. A lack of conclusive large-scale experiments, however, is a general problem in the study of complex population dynamics (Kendall et al. 1999). For ecoevolutionary models in particular, the collection of long time series of changes in phenotypes, along with data on changes in densities, could ameliorate thr problem, but such data do not yet exist, and so for now the combination of experimental and observational data that we have used to test the theory may provide the best alternative.

Moreover, the insect ecology literature suggests that eco-evolutionary dynamics may play a role in population cycles of other forest defoliating insects (Anderson and May 1981; Myers 1988, 1993). First, although previous studies used only dose-response experiments, gypsy moth dose-response experiments have shown the same trends as in our field transmission experiments (Páez et al. 2015). This in turn suggests that laboratory observations of heritable variation and costs of resistance in other insects (Boots and Begon 1993; Cory and Myers 2009; Watanabe 1987) may likewise indicate that natural selection plays a role in those insects in the field. Laboratory experiments have also detected heritable variation in responses to parasitoids in several other insect species (Kraaijeveld et al. 2002), which again may indicate effects of natural selection in the field. Finally, baculoviruses and parasitoids cause high mortality in many cycling forest defoliators (Moreau and Lucarotti 2007; Nealis 1991; Turchin 2003), and so selection pressure is often high.

The effects of seasonal breeding in our models may similarly hold for a range of host-pathogen interactions, because seasonal breeding is a widespread phenomenon in other outbreaking insects (Hunter 1991), while seasonality more generally plays a role in a range of other host-pathogen interactions (Altizer et al. 2006). More concretely, the lack of half-cycle lags in our models means that eco-evolutionary dynamics may be occurring even though half-cycle lags have not been observed. Inferring eco-evolutionary dynamics therefore appears to require not just qualitative comparisons of models to data, but also estimates of the heritability and costs of resource defenses.

Baculoviruses are also used as environmentally benign insecticides (Hunter-Fujita et al. 1998), which in the gypsy moth consists of the “Gypchek” spray product (Podgwaite et al. 1992). As is typically the case with baculoviruses of forest insects, however, Gypchek plays only a modest role in gypsy moth control, because production costs are lower for the insecticide “Btk”, a Lepidopteran-specific bacterial toxin. Because Btk targets effectively all Lepidoptera, however, concerns over its environmental costs have led to increasing public demand for Gypchek use (Boulton and Otvos 2004; Narciso 2014; Nolan 2015). Baculovirus spray products like Gypchek may therefore be used repeatedly in the future, which may alter insect outbreak cycles.

Reilly and Elderd (2014) therefore used the Dwyer et al. (2004) model to predict the longterm effects of repeated baculovirus sprays. Their work suggested that consistent spraying may dampen population cycles, eliminating outbreaks, but our eco-evolutionary models show that realistic outbreaks occur in nature for a broader range of parameters than in the Dwyer et al. model. It may therefore be the case that resistance evolution will prevent the dampening effects of repeated sprays. Extending our models to allow for repeated baculovirus sprays may thus provide a better understanding of the use of baculoviruses in microbial control, and carrying out such an extension is therefore a final important next step.

## Acknowledgments

We thank J. Armagost, P. Brandt, S. Carpenter, C. Gilroy, D. Howard, T.OHalloran, C. Maguire, Y. Ren, A. Saad, K. Smith, K. Vavra-Musser, K. Sirianni, and S. Xie, for laboratory and field assistance. We thank Viggo Andreasen and Steve Ellner for discussing our results.

## Appendix A Derivation of the Eco-Evolutionary Host-Pathogen Model

Elderd et al. (2013) present a version of the Dwyer et al. (2004) model that also includes heritable changes in average infection risk and a tradeoff between selection for reduced infection risk and selection for increased fecundity. Their model, however, assumes that the phenotypic and genotypic distributions of infection risk are identical, as though heritability *b* = 1. Here we instead assume that *b* < 1, which significantly complicates the derivation of the model.

The initial steps in the Elderd et al. derivation are nevertheless useful. First, the pathogen equation is unchanged from the original non-evolutionary insect-pathogen model of Dwyer et al. (2000), and therefore we do not consider it here. Second, by temporarily neglecting predation, we can integrate over the phenotypic distribution to derive an equation for the host population:

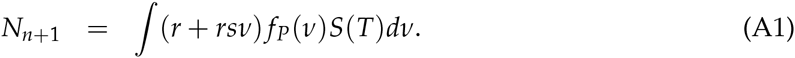

Here *fp*(*v*) is the distribution of infection-risk phenotypes *v*, and *S*(*T*) is the host density, where both are calculated after an epizootic that lasts *T* days. Host density is then calculated by allowing for both disease-driven mortality, which determines the host density after the epizootic, and by including a fecundity cost of reproduction, as determined by the cost parameters *r* and *s*. Meanwhile, Elderd et al. assumed that *T* → ∞, and used the alternative parameterization *r* + *λv* to describe the cost function, but the effects of these differences are trivial compared to the complications that arise from assuming that *b* < 1.

The key assumption in our derivation is that the epizootic reduces the mean of the distribution of phenotypes, but that it does not change the *shape* of the distribution, so that the variation parameter *V* is constant. Given the constant-shape assumption, it is possible to show that the post-epizootic mean is 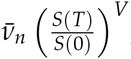, where 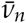 is the pre-epizootic mean (Dwyer et al. 2000). This assumption is also fundamental to the derivation of the SEIR epizootic model, equations 1-4 in the main text, which are an approximation to a model that describes the entire distribution of phenotypes. That model is highly accurate if phenotypes follow a gamma distribution, but it is only moderately inaccurate for distributions with longer tails, such as a log-normal. More importantly for our purposes, the approximation means that both the SEIR model and the multigeneration model derived here can be simulated with only modest computational costs.

To complete the derivation, we observe that the post-epizootic host density is:

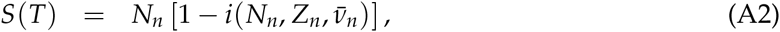

where 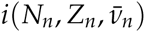 is calculated using equations 1-4. Including predation and stochasticity then gives equation 5 in the main text, which is effectively the same as the host-density equation in the Elderd et al. model.

As we described, however, the crucial difference from the Elderd et al. model is that we allow the genotypic and phenotypic distributions to have different shape parameters *V*, which is necessary to allow for imperfect heritability. This assumption becomes important when we calculate how the average phenotype 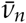 changes due to mating, because to allow for the effects of incomplete heritability, we average the fecundity costs over the *genotypic* distribution, not the *phenotypic* distribution. Specifically, we assume that the squared C.V. of the genotypic distribution is *bV*, where *V* is the squared CV of the phenotypic distribution and *b* is the heritability. Although other assumptions may also produce a reasonably simple model, this assumption has the advantage first that it ensures that genotypic variation is lower than phenotypic variation, as we would expect from quantitative genetic theory (Falconer and Mackay 1996). Also, the assumption follows an approach that is consistent with previous approaches in which quantitative genetic variation has been included in predator-prey models (Abrams and Matsuda 1997). Finally, and most importantly, the assumption produces a model that makes intuitive sense, as we will now show.

To integrate over the genotypic distribution, we proceed as follows:

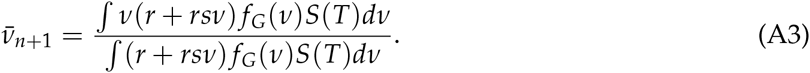

Here *f_G_*(*v*) is the post-epizootic distribution of genotypes *v*, and *S*(*T*) is again the host density after an epidemic that lasts *T* days. To solve the integral, we again use the observation that the mean of the phenotypic distribution after the epizootic is 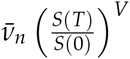. Also, we use the assumption that the genotypic distribution has the same mean as the phenotypic distribution, but that it has a squared C.V. equal to *bV*, with the proviso that, for the genotypic distribution, the mean also depends on the squared *genotypic* C.V. We then have equation 7 in the main text., which we repeat here for convenience:

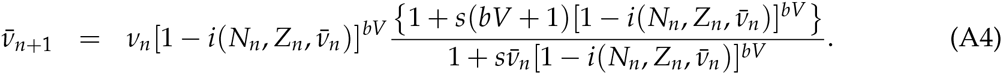

This model makes intuitive sense, in that setting *b* = 1 again produces the Elderd et al. model, while setting *b* = 0 gives the classical model with no natural selection, as in Dwyer et al. 2000 and Dwyer et al. (2004).

## Online Appendix

### Alternative models

As we mention in the main text, we considered alternative models to equations (5)-(7), as a way of testing the generality of our results. In the first of these alternatives, we eliminated stochasticity and the generalist predator, and we assumed that the epizootic model, equations (1)-(4), always proceeds to “burnout” (Keeling and Rohani 2008), which occurs if the epidemic ends because of a lack of susceptible hosts rather than because of pupation (Dwyer et al. 2000). Strictly speaking, the burnout approximation requires the assumption that *t* → ∞, but in spite of this seemingly unrealistic assumption, the burnout approximation’s prediction of the fraction infected is often close to the prediction of equations (1)-(4) with a realistic epizootic length of 8 weeks (Fuller et al. 2012). Allowing epizootics to end because of pupation instead of burnout is quantitatively important when we attempt to reproduce outbreak cycles, because when epizootics are only 8 weeks long, the model produces longer period, larger amplitude cycles that better match the outbreak data (Fuller et al. 2012). Including a model that allows for the burnout approximation nevertheless allows us to relax the assumption that epizootics are ended by pupation, and it allows us to use qualitative stability analysis, which permits a deeper understanding of the effects of natural selection on population cycles.

Given these simplifications, equations (5)-(7) in the main text become:

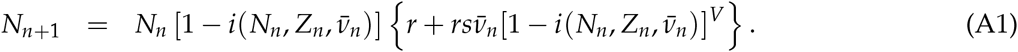

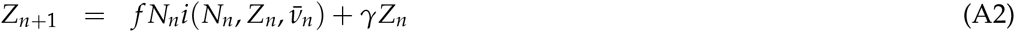

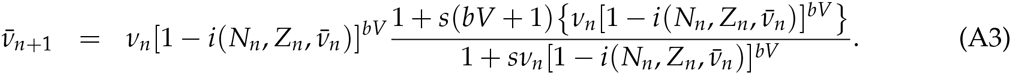

Here the fraction of infected individuals, *i*, is represented by the burnout approximation, which is calculated from equations (1)-(4) by allowing *t* → ∞. In that case, we can write down an implicit expression for *i*:

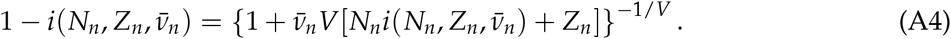

Using this expression in equations (A1)-(A3) produces a model for which we can easily carry out stability analysis.

The initial step is to calculate the model equilibria. For this model, we cannot prove that there is only one non-zero equilibrium, but two lines of indirect evidence suggest that multiple equilibria are unlikely unless baseline fecundity *r* > 1. First, if we set **b** = 1, equations (A1)-(A4) are the same as the no-predator model in Elderd et al. (2008), for which it is possible to prove that *r* < 1 guarantees that there is only one equilibrium. We have not been able to prove a similar result for equations (A1)-(A4), but numerical iteration of the model suggests that multiple equilibria are at least unlikely unless *r* is quite close to 1, as we will show. These considerations are important because a lack of multiple equilibria makes calculation of the Jacobian matrix reasonably straightforward. To write down the Jacobian, we first rescale the model, by defining:

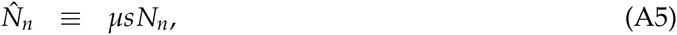

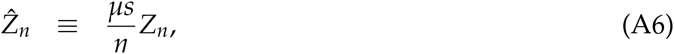

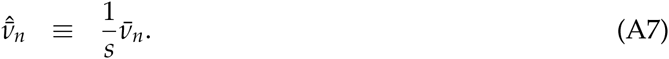

For convenience, we also define 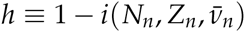, to produce:

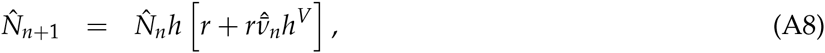

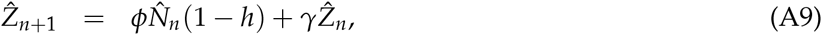

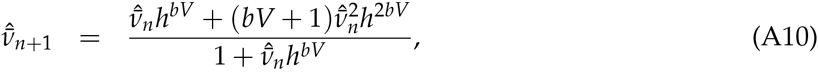

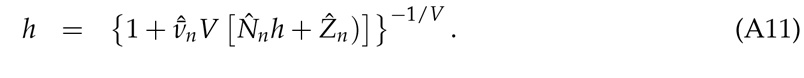

We can then use equations (A8)-(A11) to find the equilibrium conditions, such that 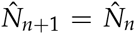, 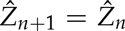, and 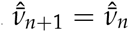. Also for convenience, we drop the^ symbols, and use *’s to label the equilibrium values of the state variables:

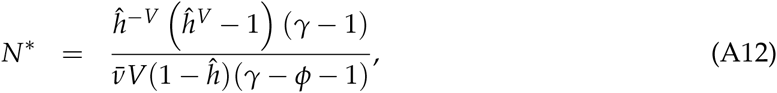

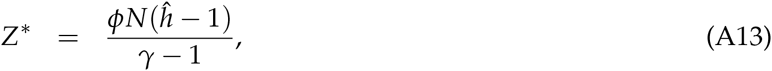

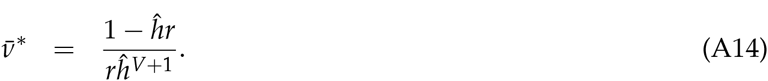

Here 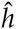 is the the fraction of individuals uninfected at equilibrium, such that the fraction infected is calculated from equation (A4). By replacing 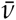 in equation (A10) by 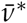 (equation (A14)), we can write down an implicit expression for *h*, which we can easily solve numerically:

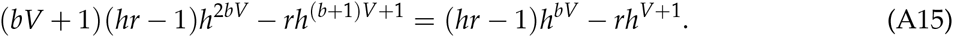

Plotting the left-hand and right-hand sides of this equation for a range of parameter values suggests that, for *r* < 1, there is only one internal equilibrium, but for *r* > 1, two equilibria often occur, with dynamics that generally lead to the extinction of the pathogen population. Given that realistic values of *r* are all well below 1, in what follows we concentrate on the case for which *r* < 1.

Next, we define 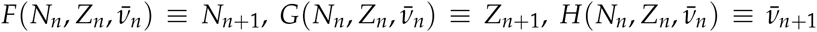 and we differentiate each function with respect to *N_n_, Z_n_* and 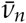, to produce the Jacobian matrix. To do this, we first differentiate *h* with respect to *N, Z* and 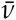.

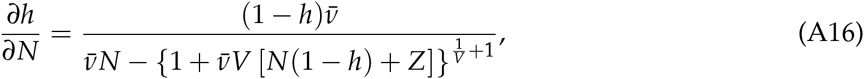

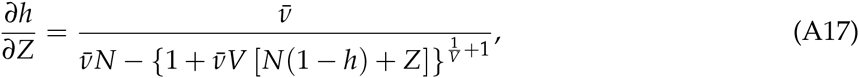

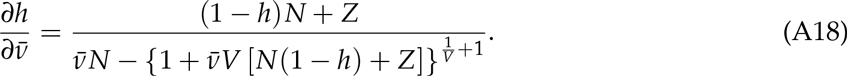

The matrix of partial derivatives is then described by the matrix *J* above.

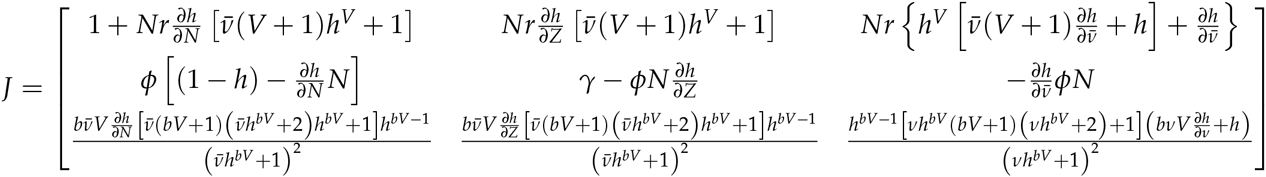

By numerically solving equation (A15) for the equilibrium value 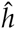, we can calculate the equilibrium values of the state variables *N*\*, *Z*\* and 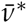, and insert them into the matrix *J* to construct the Jacobian matrix. We can then numerically solve for values of the parameters for which the Jacobian has one real eigenvalue and a complex pair of eigenvalues with modulus 1, the criterion that defines the boundary between a stable point equilibrium and a Hopf bifurcation, and thus population cycles. In Fig. A1, the shading denotes periods and amplitudes as calculated by numerical iteration of equations (A8)-(A11), while the unshaded region is the area within which only a stable point equilibrium occurred. The dashed line in Fig. A1 then shows that the eigenvalue prediction of the boundary between cycles and a stable equilibrium point matches the boundary between cycles and a stable equilibrium from the numerical iterations, confirming both calculations. At higher heritability, however, the stability calculation breaks down for very high values of the fecundity cost *r*. In that region, it appears that the model may show multiple equilibria, with the proviso that such high values of *r* are unlikely to be biologically realistic and are therefore of limited interest.

**Figure A1:**
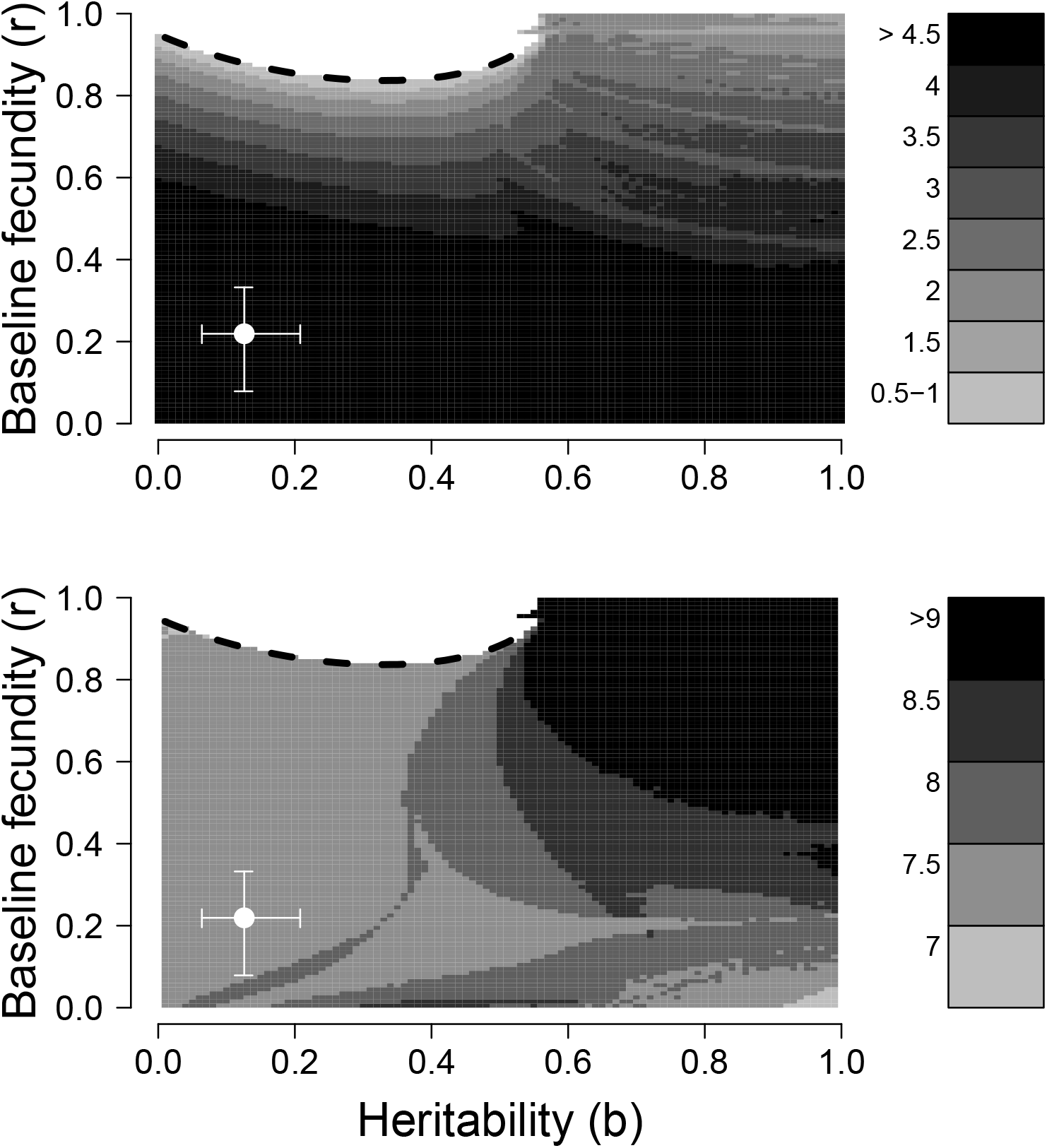
The average amplitude (top) and period (bottom) of population cycles in equations (A1)-(A4) for different combinations of the heritability of infection risk, *b*, and baseline reproduction, *r*. The white space at high baseline fecundity and low to moderate heritability indicates regions in which a stable equilibrium point occurs.

More generally, Fig. A1 shows that for equations (A1)-(A3), increasing values of *r* are generally stabilizing, as in equations (5)-(7) in the main text. In contrast to equations (5)-(7), however, for this model low values of heritability are no longer destabilizing, most likely because equations (A1)-(A3) assume that epizootics are terminated by burnout rather than by pupation. Pupation tends to terminate epizootics sooner than burnout would, exacerbating the effects of the delayed density-dependence that drives cycles (Fuller et al. 2012), an effect that is apparently stronger when heritability is low. The result is that the long periods and larger amplitudes seen at low heritability in fig. 3 in the main text are eliminated, as is the area of dramatic oscillations at high heritability and intermediate cost.

We also considered a host-parasitoid model, in which there is only one parasitoid generation per host generation, as compared to the multiple pathogen generations per host generation in equations (1)-(4). As in equations (1)-(4), however, we again allow for host variation, this time in the risk of being successfully attacked by the parasitoid. We then have a closed-form expression for the fraction infected (Godfray 1994):

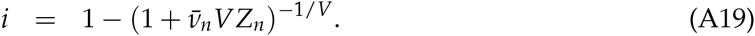

We then again add a generalist predator, on the grounds that high levels of generalist predator attacks have been observed not just in defoliators whose outbreaks are driven by pathogens, but also in defoliators in which outbreaks are instead driven by parasitoids (Dwyer et al. 2004). The model then differs from the multi-generation model in the text, equations (5)-(7), only in using equation (A19) to calculate the fraction infected 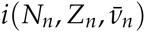.

Fig. A2 shows a time series of the host-parasitoid model dynamics, to show that consumer-resource cycles are again partly driven by fluctuations in the average risk of attack, as in equations (5)-(7) in the main text. Also, Fig. A3 shows the average period and the average amplitude for this model for a range of parameters, as in Fig. 3 in the main text. The effects of the parameters on the host-parasitoid-predator model are thus qualitatively similar to the effects of the parameters on the host-pathogen-predator model, with the proviso that, for values of heritability much above 0.4, the parasitoid goes extinct unless the cost parameter *r* is close to 1. This instability likely occurs because the parasitoid has only one generation per year, which exacerbates the effects of the delayed density-dependence that drives cycles, and because of the lack of a parasitoid functional response, which we have omitted to allow for a more straightforward comparison to the model in the main text.

**Figure A2:**
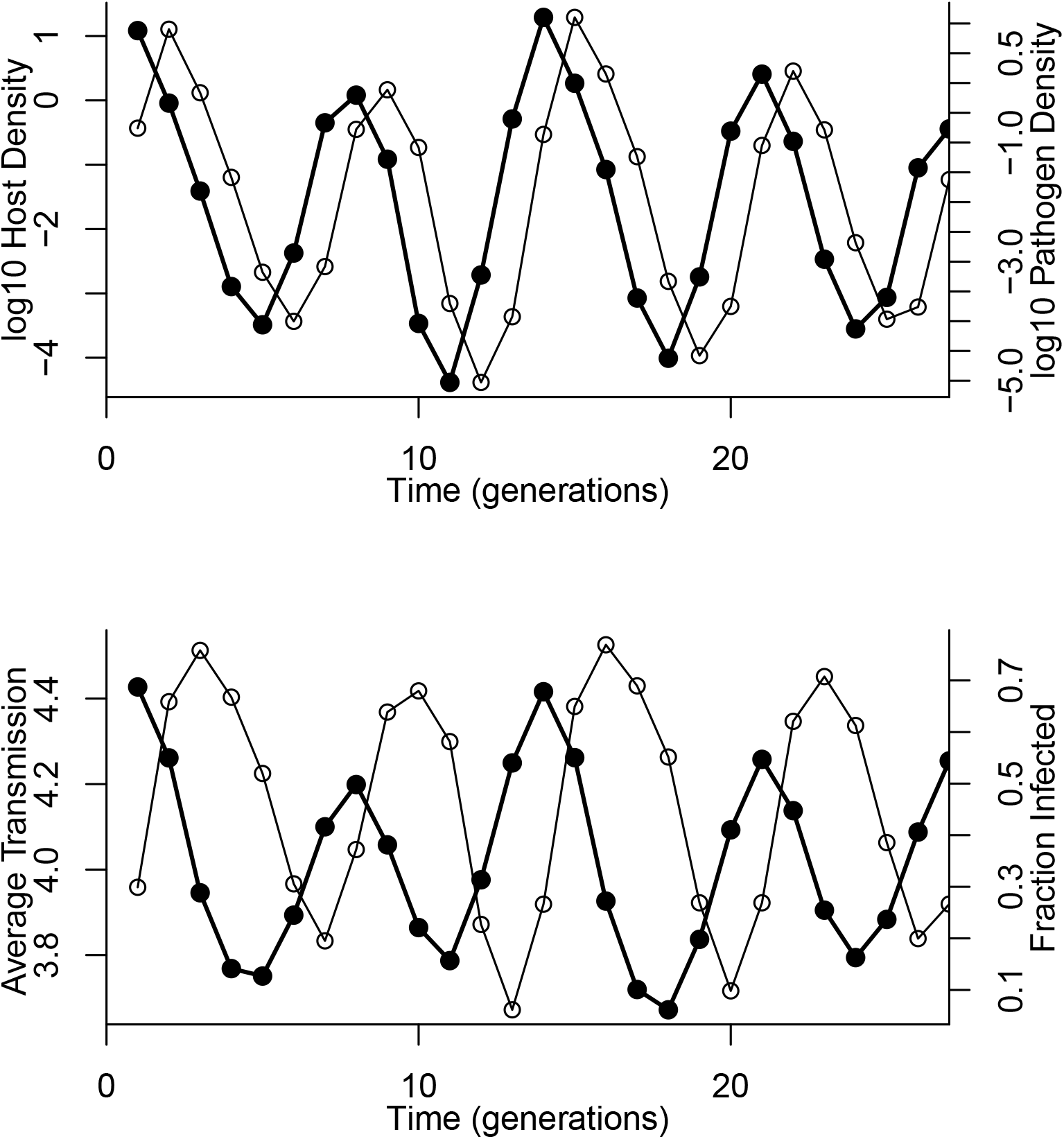
Predictions of an eco-evolutionary host-parasitoid model. As in Fig. 1 in the main text, the top panel shows changes in host and parasitoid density (black and grey lines, respectively), while the bottom panel shows the corresponding changes in the parasitoid attack rate and the fraction infected (also black and grey lines, respectively). Parameter values are *b* = 0.13, *r* = 0.42, *s* = 0.5, *V* = 10, *γ* = 0.3, *ϕ* = 1.0, *a* = 0.96, *ω* = 0.14.

**Figure A3:**
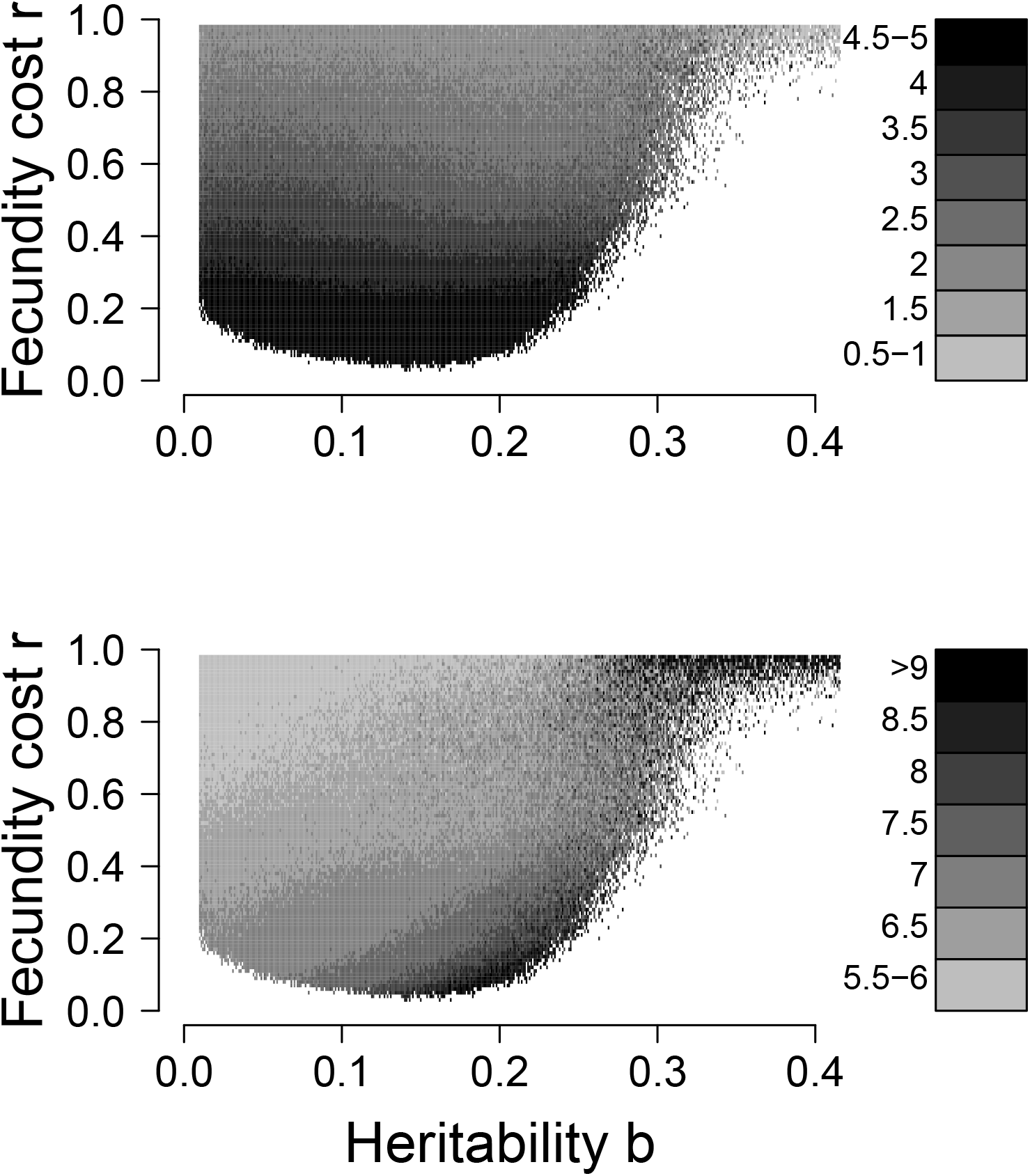
Effects of baseline fecundity *r* and heritability *b* on the period and amplitude of outbreak cycles in the eco-evolutionary host-parasitoid model. The top panel shows average cycle amplitudes, in orders of magnitude, while the bottom panel shows the average period, in years.

### Details of insect rearing methods

Wild egg masses were collected from 5 different sites in the Midwestern states of the U.S (Páez et al. 2015). Previous work has shown that geographic structure in infection risk across gypsy moth populations is slight (Elderd et al. 2008), and previous dose-response experiments showed that these particular populations had very similar responses in laboratory dose-response experiments (Fleming-Davies et al. 2015). It therefore appeared that the initial populations would have similar infection risk in the field.

Before hatch, we soaked all egg masses in a 4% formalin solution, which surface sterilizes the eggs by killing any occlusion bodies on their surface. This procedure kept virus mortality in control treatments low, as it has been in previous experiments (Elderd et al. 2008; Fuller et al. 2012). Hatching larvae were then reared in groups of 30, in 177 ml (6 oz) plastic cups, containing 50-100 ml of artificial diet, at 25 °C in an incubator with a 14:10 light-dark cycle, following long established rearing procedures for this insect (McManus and Doane 1981).

To produce larvae for experiments, we reared insects from wild-collected egg masses to adulthood, and we mated the adults. Matings were usually conducted within 24 hours of female emergence. Full siblings can be easily produced by exposing one male to one female. To produce half siblings, we instead exposed one male to 2-3 virgin females every 24 hours (Páez et al. 2015). After reproduction and egg mass deposition, we allowed 28 days of pre-diapause at 25 °C, and then we induced diapause by cooling the egg masses to 5° C for 9-10 months. The most important rounds of transmission in nature occur when larvae are in the fourth instar (Woods and Elkinton 1987), and so we used fourth instars in all of our experiments.

Infection risk given virus exposure can vary within a larval stage or instar (Grove and Hoover 2007), so we synchronized the uninfected insects before deployment in the field. To do this, we collected larvae in the third instar whose head capsules had slipped forward, indicating that they were within 24 hours of molting to the next instar. We held these insects at 5° C in the lab until we had enough insects to begin an experiment, which typically took 48 hours. This procedure ensures that the measurement error in field transmission experiments is no higher than what we would expect from binomial sampling (Elderd et al. 2008).

### Calculating Fecundity Costs Including Lab-reared Insects, and Allowing for Density-Independent Mortality

As we mentioned in the main text, in estimating the cost of resistance, we included only insects that had survived virus exposure in the field, but in a second analysis we included insects reared in the lab, where larvae could not encounter the virus. Here we present the results from this second analysis. As in the first analysis, we used a linear mixed effects model in which pupal weight was a function of infection risk *ν* and random family effects, but we also included the effects of the rearing method, meaning lab versus field. Again as we mentioned in the main text, AIC analysis of regression models showed that the best model for the combined data set included different intercepts, reflecting the higher pupal weights associated with consumption of artificial diet, but there was nevertheless a common slope for field-reared and lab-reared insects (Table A1, Fig. A4). The fecundity cost of reduced infection risk was thus indistinguishable for field-reared and lab-reared insects.

**Figure A4:**
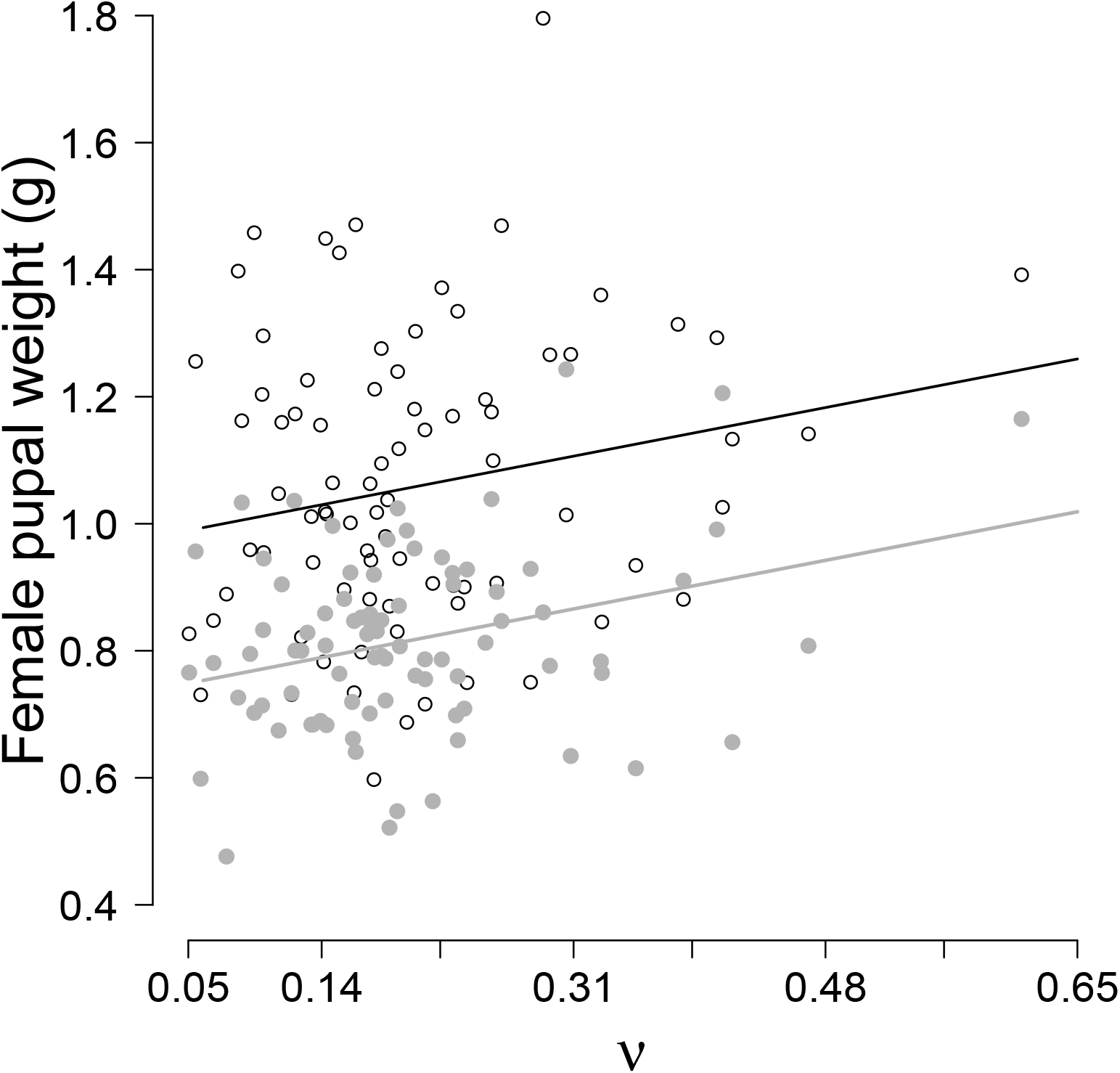
Relationship between infection risk *ν* and female pupal weight, including both insects that survived the field experiment (closed grey circles and grey line), and unexposed insects that were only reared in the lab (open black circles and black line). Lines are based on median values for the average model coefficients from a bootstrapping procedure, as described in the text. The higher average weights of lab-reared insects result from the use of artificial diet in the lab.

**Table A1:** AIC analysis for models fit to data for both lab-reared and field-reared insects. “Intercepts differ” means that the intercepts were different for the two groups, and likewise for the slopes.

| # | Model | ΔAIC | AIC Weight |
| --- | --- | --- | --- |
| 1 | **Intercepts differ, slopes same** | **0** | 0.88 |
| 2 | Intercepts differ, slopes differ | 1.97 | 0.12 |
| 3 | Intercepts differ, slopes = 0 | 12.6 | $< 10^{-5}$ |
| 4 | Intercepts and slopes same | 54.6 | $< 10^{-5}$ |
| 5 | Intercepts same, slopes = 0 | 63.2 | $< 10^{-5}$ |

To estimate the uncertainty in the parameters of the regression lines, we bootstrapped the best model ten thousand times, such that, at each iteration, the model was re-fit using re-sampled values of pupal weights and infection risks *ν*. These results showed that the median intercept for field-reared insects was 0.73 (upper and lower 95th percentiles = 0.70,0.80); the median (common) slope was 0.45 (upper and lower 95th percentiles = 0.09, 0.56); and the median intercept for laboratory-reared insects was 0.97 (upper and lower 95th percentiles = 0.93, 1.05). The lower confidence bound on the slope parameter does not include 0, and we therefore conclude that including lab-reared insects does not affect our conclusion that there is a significant relationship between infection risk *ν* and pupal weight.

To estimate the cost parameters *r* and *s*, we first converted from pupal weight to egg mass weight using data from Páez et al. (Páez et al. 2015), and second from egg mass weight to egg number using data from Dwyer and Elkinton (1995). An additional consideration, however, is that the model requires an estimate of net fecundity, because hatchling gypsy moth larvae die from many causes, including starvation during dispersal from egg masses, which are laid on bark, to foliage at the ends of branches. To allow for such losses, we used data from Hunter and Elkinton (2000), who measured early instar survival in experimental gypsy moth populations. To account for the overall uncertainty in these conversions, we bootstrapped the relevant data, and we used the resulting resampled model coefficients to calculate *r* and *s*. Our estimates for field F and lab L insects were then: *S_F_* = 1.21 (0.2,1.9), *r_F_ =* 0.22 (0.05,0.51) and *S_L_* = 0.74 (0.13,1.03), *r_L_* = 0.36 (0.08,0.76). As we mention in the main text, in our model we used only estimates for field-reared insects.

### Details of statistical analyses

Because over-dispersion levels in field transmission experiments are generally low (Elderd et al. 2008), in our likelihood function, we described the chance of infection using a binomial distribution (McCullagh and Nelder 1989). Also, standard quantitative genetic practice is to assume that the effects due to sire, dam and experimental day follow normal distributions with mean 0 (Falconer and Mackay 1996). Our likelihood function is therefore:

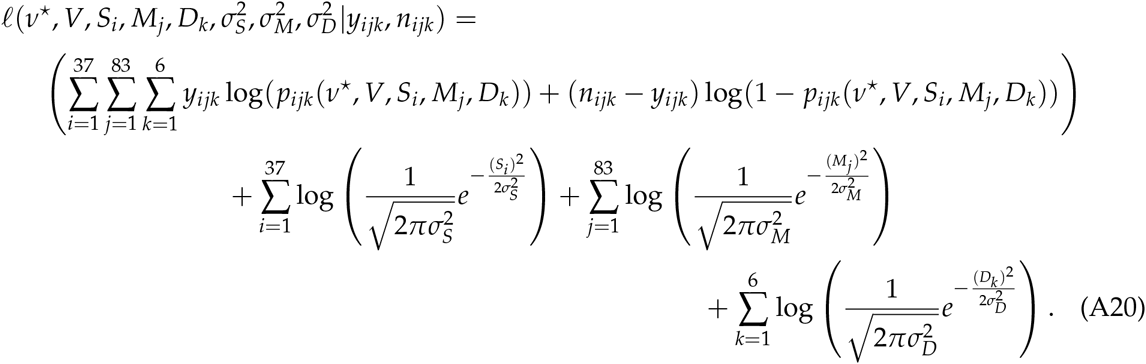

Here 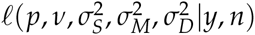 is the log-likelihood of the transmission parameters ν^*^ and V, the sire effect *S_i_*, the maternal effect *M_j_*, the day effect *D_k_*, and the corresponding variances 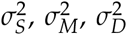, given the the number infected *y_ijk_* and the total sample size *n_ijk_*. Also, *p_ijk_ (ν_*_, V, S_i_, M_j_, D_k_*) is the prediction of the transmission equation (8) in the main text for insects with sire *i*, dam *j*, and start-day *k*, given baseline infection risk *ν^*^* and squared coefficient of variation *V*. The bounds on the summations reflect the number of treatments, such that there were 37 sires, 83 dams, and 6 start days. Estimating heritability therefore required that we estimate 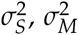 and 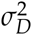, which in turn required that we simultaneously estimate the random effect sizes *S_i_*, *M_j_* and *D_k_*, as well as the baseline infection risk *ν^*^* and the squared coefficient of variation *V*.

To do this, we used a Bayesian hierarchical model, in combination with a Metropolis-Hastings MCMC algorithm. Specifically, we used 10 Markov-chain Monte Carlo chains that were sampled every 1000th step over 1.5 x 10^6^ steps after discarding the first 1.5 x 10^4^ steps. Chain convergence was confirmed using the Gelman-Rubin diagnostic criterion (Plummer et al. 2006). To calculate heritability, we then inserted the sets of values of 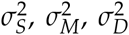, *σD* from the posterior distribution of the parameters into the heritability equation (10) in the main text.

### Effects of variation in cost-scaling *s* and the long-term pathogen survival *γ* on the model predictions

Fig. A5 shows that increasing the value of *s* to the upper bound on its 95% HPD leads to periods and amplitudes that are almost identical to the periods and amplitudes seen when the costscaling parameter *s* = 1.21, the median value of *s*, as shown by comparison to Fig. 3 in the main text. The major difference between the two cases is that, when *s* = 1.9, the large-amplitude, long-period fluctuations that occur at high heritability for *s* = 1.21 are almost eliminated. Meanwhile, Fig. A6 shows that reducing *s*, so that *s* = 0.2, the lower bound on the 95% HPD of *s*, gives periods and amplitudes that are only modestly larger, at least for realistic values of heritability b and baseline fecundity *r*. When heritability is higher, however, the wild fluctuations that occur at intermediate fecundity cost *r* are so dramatic that the host and/or the pathogen often goes extinct (white space in Fig. A6). Reductions in the cost parameter *s* are thus mildly destabilizing, unless heritability is very high, basically for the same reasons that reductions in *r* are destabilizing.

**Figure A5:**
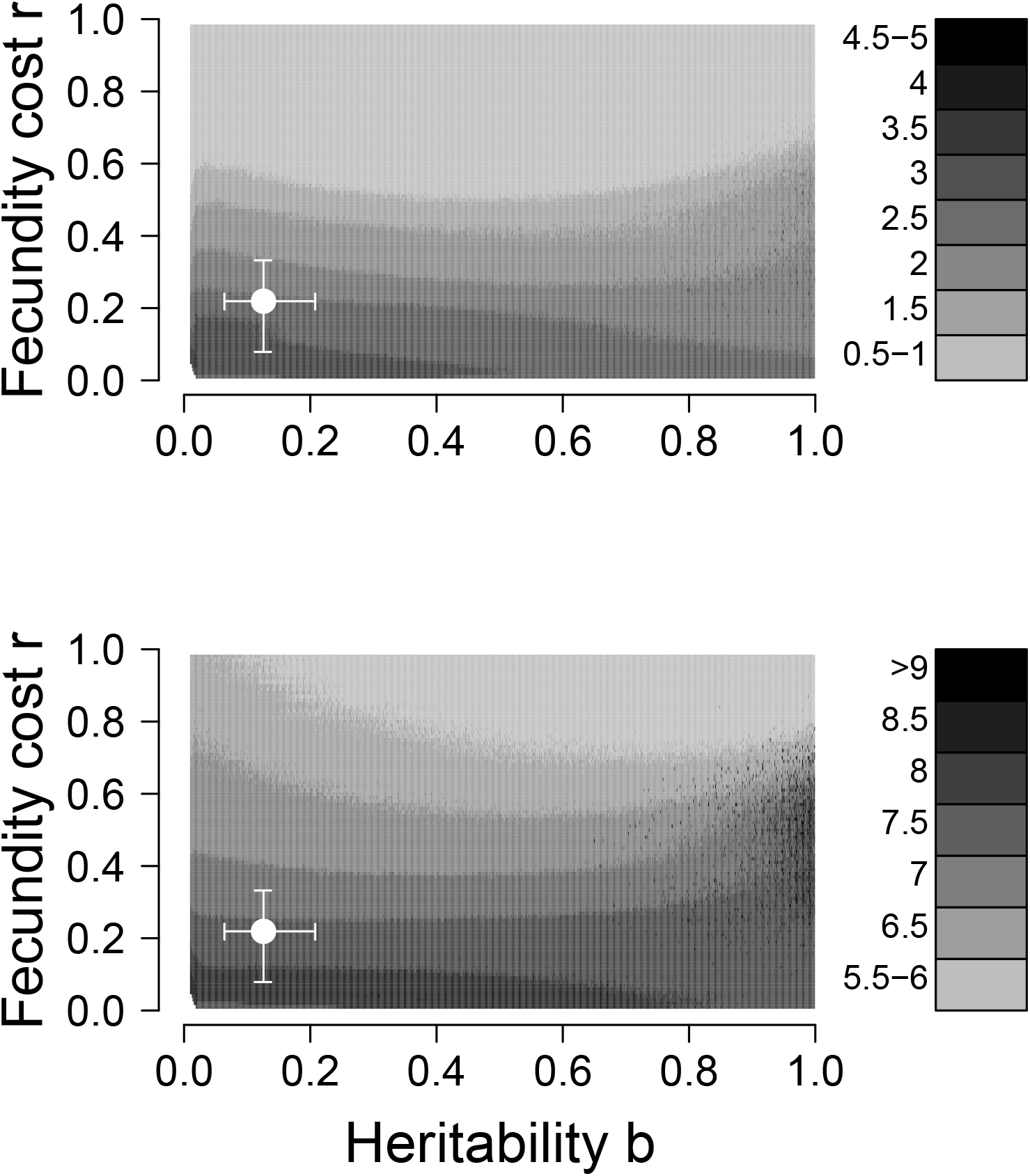
Effects of baseline fecundity *r* and heritability *b* on the period and amplitude of outbreak cycles in the long term model, equations (5)-(7), as in Fig. 3 in the main text, except that here the cost-scaling parameter *s* = 1.9. Again the top panel shows average cycle amplitudes in orders of magnitude, while the bottom panel shows the average period in years.

**Figure A6:**
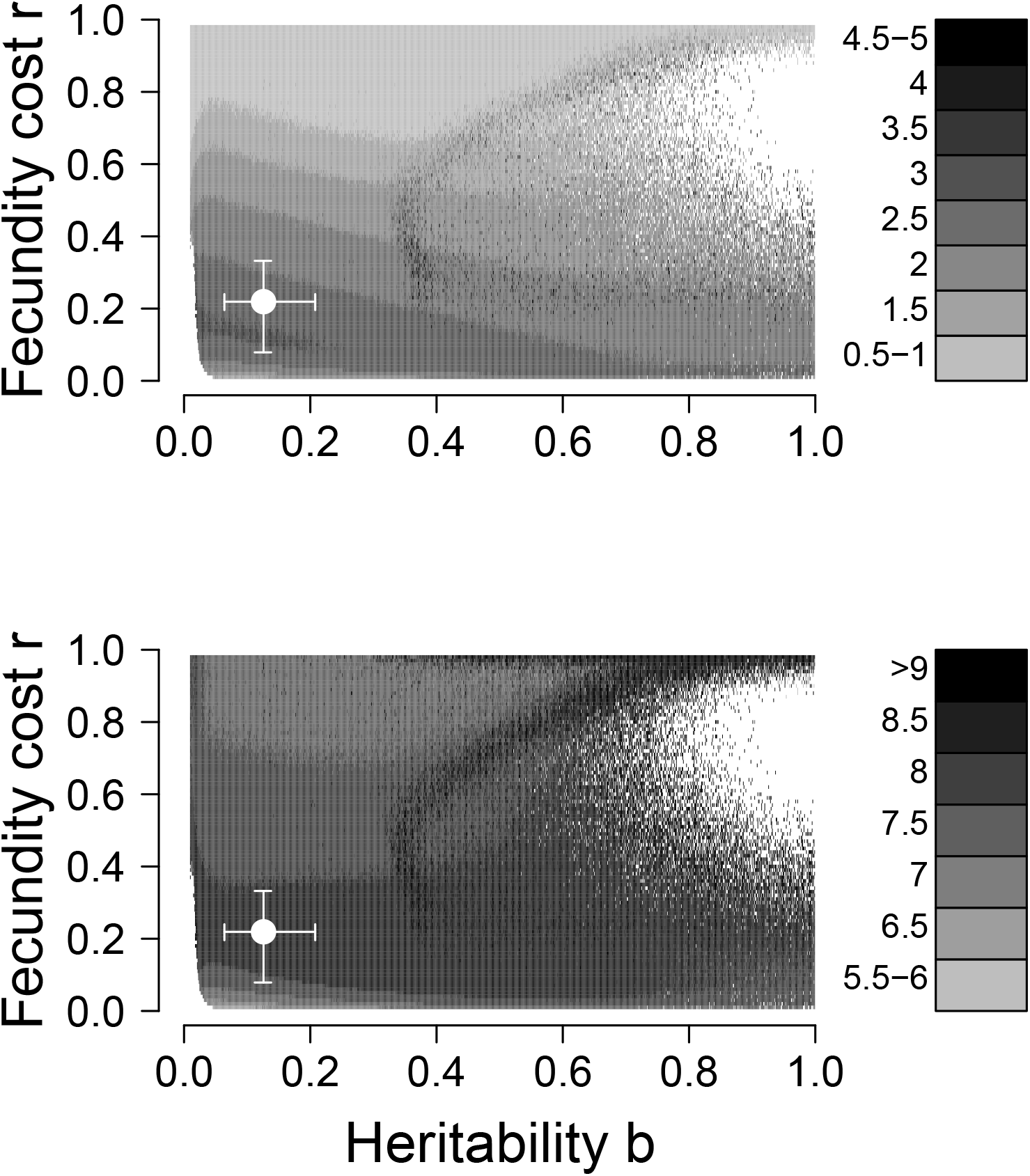
As in Fig. 3, except that the cost-scaling parameter *s* = 0.2.

Reductions in the long-term pathogen survival parameter *γ* also have complex effects. As Fig. A7 shows, simply reducing *γ* leads to shorter periods and smaller amplitudes, although both periods and amplitudes again fall in a realistic range. This effect likely occurs because lower pathogen survival leads to reduced selection intensity, and thus less violent fluctuations in population densities. Reduced long-term survival, however, also increases the chance that the pathogen will go extinct, increasing the size of the region at high heritability for which the model simply crashes.

**Figure A7:**
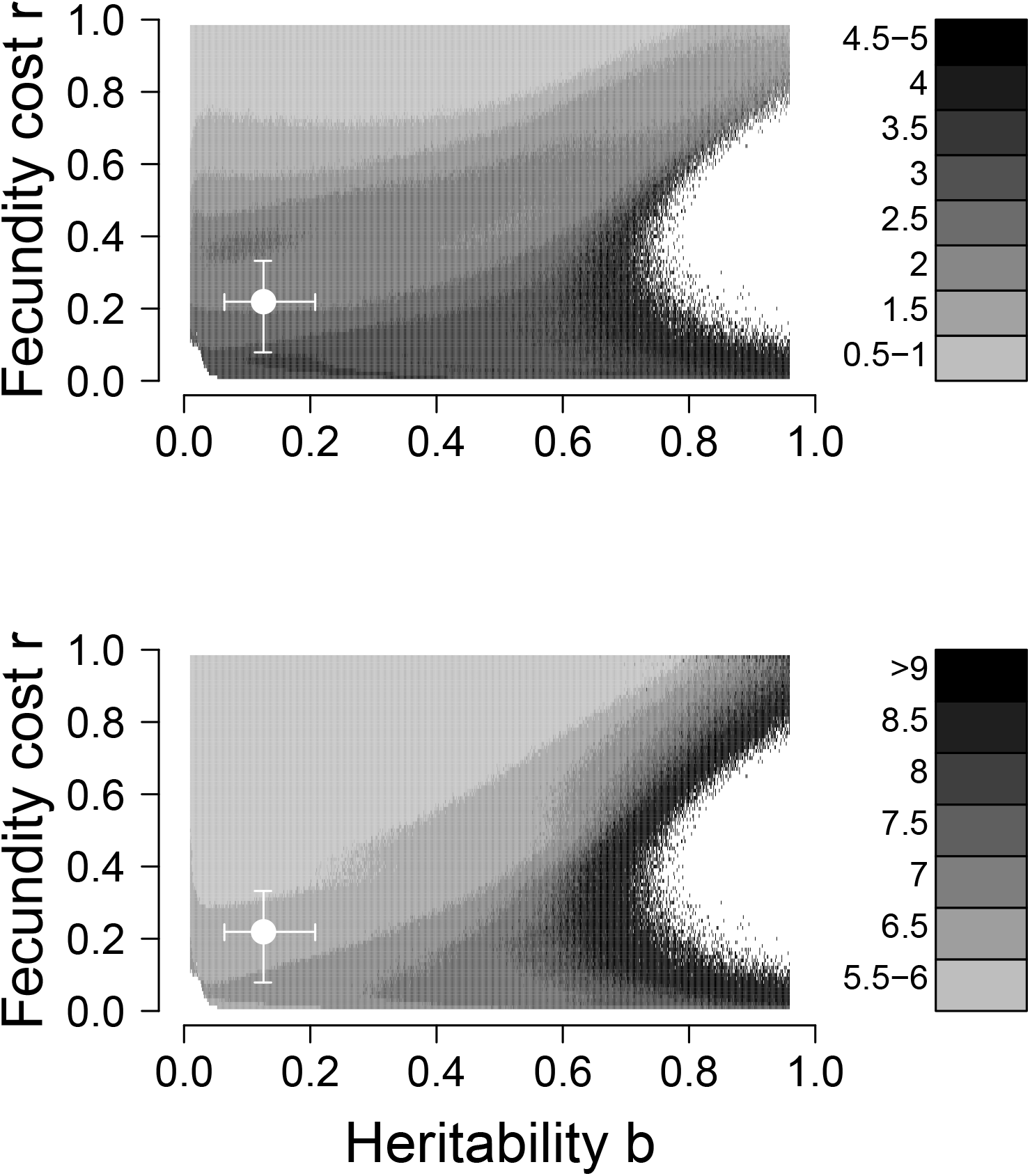
As in Fig. 3, except that the long-term pathogen survival parameter *γ* = 0.1.

Fig. A8 again shows periods and amplitudes when *γ* = 0.1, but for values of the cost-scaling parameter *s* = 0.2, the lower bound on the 95% HPD. Here *γ* has similar effects to when *s* = 1.21, except that periods are longer and amplitudes are larger, and the region of pathogen extinction is also larger. Given that reductions in cost-scaling *s* have similar effects to reductions in fecundity costs *r*, both effects are what we would expect for reduced values of *s*.

**Figure A8:**
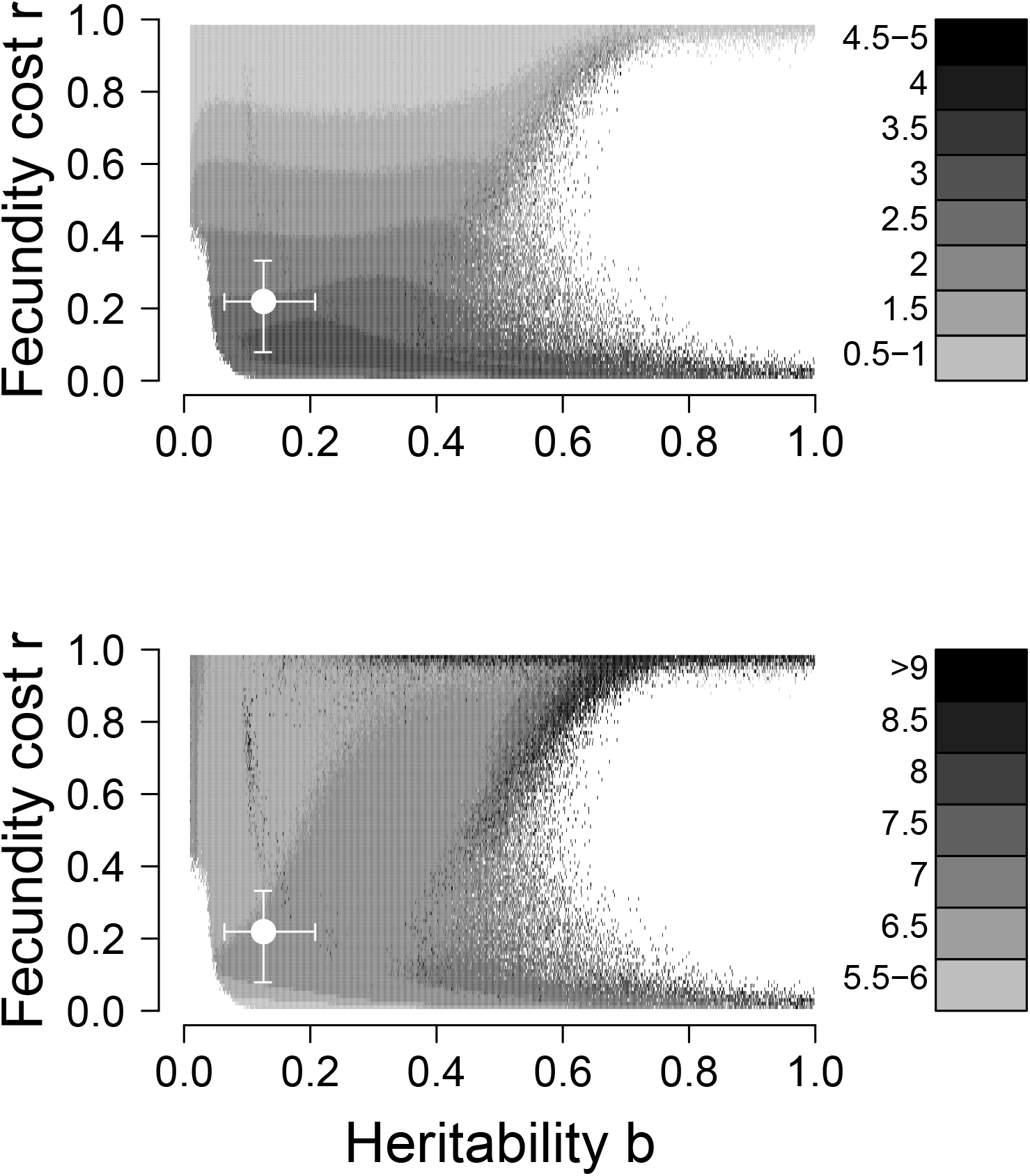
As in Fig. 3, except that the cost-scaling parameter *s* = 0.2, and the long-term pathogen survival parameter *γ* = 0.1.

## References

Abrams, P. 2000. The evolution of predator-prey interactions: Theory and evidence. Annu. Rev. Ecol. Evol. Syst. 31:79–105.

Abrams, P. A., and H. Matsuda. 1997. Prey adaptation as a cause of predator-prey cycles. Evolution 51:pp. 1742–1750.

Allstadt, A. J., K. J. Haynes, A. M. Liebhold, and D. M. Johnson. 2013. Long-term shifts in the cyclicity of outbreaks of a forest-defoliating insect. Oecologia 172:141–151.

Altizer, S., A. Dobson, P. Hosseini, P. Hudson, M. Pascual, and P. Rohani. 2006. Seasonality and the dynamics of infectious diseases. Ecol. Lett. 9:467–484.

Altizer, S., D. Harvell, and E. Friedle. 2003. Rapid evolutionary dynamics and disease threats to biodiversity. Trends in Ecology & Evolution 18:589–596.

Anderson, R. M., and R. M. May. 1981. The population dynamics of micro-parasites and their invetebrate hosts. Phil. Trans. Roy. Soc. B. 291:451–524.

Auld, S. K. J. R., S. R. Hall, J. H. Ochs, M. Sebastian, and M. A. Duffy. 2014. Predators and patterns of within-host growth can mediate both among-host competition and evolution of transmission potential of parasites. Am. Nat. 184:S77–S90.

Auld, S. K. J. R., R. M. Penczykowski, J. H. Ochs, D. C. Grippi, S. R. Hall, and M. A. Duffy. 2013. Variation in costs of parasite resistance among natural host populations. J. Evol. Biol. 26:2479–2486.

Bjørnstad, O. N., C. Robinet, and A. M. Liebhold. 2010. Geographic variation in North American gypsy moth cycles: subharmonics, generalist predators, and spatial coupling. Ecology 91:106–118.

Boots, M., and M. Begon. 1993. Tradeoffs with resistance to a granulosis virus in the Indian meal moth examined by a laboratory evolution experiment. Functional Ecology 7:528–534.

Boulton, T., and I. Otvos. 2004. Monitoring native non-target Lepidoptera for three years following a high dose and volume application of Bacillus thuringiensis subsp kurstaki. Int. J. Pest Manag. 50:297–305.

Box, G. 1979. Robustness in Statistics, chap. Robustness in the strategy of scientific model building, pages 202–236. Academic Press, N.Y.

Capinera, J. L., K. P. Kirouac, and P. Barbosa. 1976. Phagodeterrency of cadaver components to gypsy moth larvae, Lymantria dispar. J. Invertebr. Pathol. 28:277–279.

Cory, J. S., and K. Hoover. 2006. Plant-mediated effects in insect-pathogen interactions. Trends Ecol. Evol. 21:278–286.

Cory, J. S., and J. H. Myers. 2009. Within and between population variation in disease resistance in cyclic populations of western tent caterpillars: a test of the disease defence hypothesis. J. Anim. Ecol. 78:646–655.

Dieckmann, U. 2002. Adaptive dynamics of pathogen-host interactions. Chap. 4, pages 39–59 in U. Dieckmann, K. Sigmund, and H. Metz, eds. Adaptive Dynamics of Infectious Diseases: In Pursuit of Virulence Management. Cambridge University Press, Cambridge, U.K.

Diss, A., J. Kunkel, M. Montgomery, and D. Leonard. 1996. Effects of maternal nutrition and egg provisioning on parameters of larval hatch, survival and dispersal in the gypsy moth, Lymantria dispar L. Oecologia 106:470–477.

Doebeli, M. 1997. Genetic variation and persistence of predator-prey interactions in the Nicholson-Bailey model. J. Theor. Biol. 188:109–120.

Dwyer, G., J. Dushoff, J. Elkinton, and S. Levin. 2000. Pathogen-driven outbreaks in forest defoliators revisited: Building models from experimental data. Am. Nat. 156:105–120.

Dwyer, G., J. Dushoff, J. S. Elkinton, J. P. Burand, and L. S. A. 2002. Variation in susceptibility: lessons from an insect virus. Chap. 6, pages 74–84 in K. S. U. Dieckmann and H. Metz, eds. Adaptive Dynamics of Infectious Diseases: In Pursuit of Virulence Management. Cambridge University Press, Cambridge, U.K.

Dwyer, G., J. Dushoff, and S. H. Yee. 2004. The combined effects of pathogens and predators on insect outbreaks. Nature 430:341–345.

Dwyer, G., J. Elkinton, and J. Buonaccorsi. 1997. Host heterogeneity in susceptibility and disease dynamics: Tests of a mathematical model. Am. Nat. 150:685–707.

Dwyer, G., and J. S. Elkinton. 1993. Using simple models to predict virus epizootics in gypsy moth populations. J. Anim. Ecol. 62:1–11.

Dwyer, G., and J. S. Elkinton. 1995. Host dispersal and the spatial spread of insect pathogens. Ecology 76:1262–1275.

Dwyer, G., J. Firestone, and T. Stevens. 2005. Should models of disease dynamics in herbivorous insects include the effects of variability in host-plant foliage quality? Am. Nat. 165:16–31.

Eakin, L., M. Wang, and G. Dwyer. 2015. The effects of the avoidance of infectious hosts on infection risk in an insect-pathogen interaction. Am. Nat. 185:pp.100–112.

Elderd, B. D. 2013. Developing models of disease transmission: Insights from ecological studies of insects and their baculoviruses. PLoS Pathogens 9.

Elderd, B. D., J. Dushoff, and G. Dwyer. 2008. Host-pathogen interactions, insect outbreaks, and natural selection for disease resistance. Am. Nat. 172:829–842.

Elderd, B. D., B. J. Rehill, K. J. Haynes, and G. Dwyer. 2013. Induced plant defenses, host-pathogen interactions, and forest insect outbreaks. Proc. Natl. Acad. Sci. 110:14978–14983.

Elkinton, J. S., and A. M. Liebhold. 1990. Population dynamics of gypsy moth in North America. Annual Review of Entomology 35:571–596.

Ellner, S. P. 2013. Rapid evolution: from genes to communities, and back again? Functional Ecology 27:1087–1099.

Ellner, S. P., M. A. Geber, and N. G. Hairston. 2011. Does rapid evolution matter? measuring the rate of contemporary evolution and its impacts on ecological dynamics. Ecol. Lett. 14:603–614.

Falconer, D., and T. Mackay. 1996. Introduction to quantitative genetics. Prentice Hall, Harlow, England.

Fleming-Davies, A. E., V. Dukic, V. Andreasen, and G. Dwyer. 2015. Effects of host heterogeneity on pathogen diversity and evolution. Ecology Letters 18:1252–1261.

Fleming-Davies, A. E., and G. Dwyer. 2015. Phenotypic variation in overwinter environmental transmission of a baculovirus and the cost of virulence. The American Naturalist 186:797–806.

Fuller, E., B. D. Elderd, and G. Dwyer. 2012. Pathogen persistence in the environment and insect-baculovirus interactions: Disease-density thresholds, epidemic burnout, and insect outbreaks. Am. Nat. 179:pp.E70–E96.

Fussmann, G. F., S. P. Ellner, K. W. Shertzer, and N. G. Hairston Jr. 2000. Crossing the hopf bifurcation in a live predator-prey system. Science 290:1358–1360.

Godfray, H. C. J. 1994. Parasitoids : behavioral and evolutionary ecology. Princeton University Press.

Grove, M. J., and K. Hoover. 2007. Intrastadial developmental resistance of third instar gypsy moths (Lymantria dispare L.) to L. dispar nucleopolyhedrovirus. Biol. Control 40:355–361.

Hajek, A. E. 1999. Pathology and epizootiology of Entomophaga maimaiga infections in forest Lepidoptera. Microbiol. Mol. Biol. Rev. 63:814–835.

Hajek, A. E., P. C. Tobin, and K. J. Haynes. 2015. Replacement of a dominant viral pathogen by a fungal pathogen does not alter the collapse of a regional forest insect outbreak. Oecologia 177:785–797.

Haynes, K. J., A. M. Liebhold, T. M. Fearer, G. Wang, G. W. Norman, and D. M. Johnson. 2009. Spatial synchrony propagates through a forest food web via consumer-resource interactions. Ecology 90:2974–2983.

Haynes, K. J., A. M. Liebhold, and D. M. Johnson. 2009. Spatial analysis of harmonic oscillation of gypsy moth outbreak intensity. Oecologia 159:249–256.

Henter, H., and S. Via. 1995. The potential for coevolution in a host-parasitoid system. 1. The potential genetic variation within an aphid population in susceptibility to a parasitic wasp. Evolution 49:427–438.

Herzog, J., C. B. Mueller, and C. Vorburger. 2007. Strong parasitoid-mediated selection in experimental populations of aphids. Biol. Lett. 3:667–669.

Hiltunen, T., N. G. Hairston, Jr., G. Hooker, L. E. Jones, and S. P. Ellner. 2014. A newly discovered role of evolution in previously published consumer-resource dynamics. Ecol. Lett. 17:915–923.

Hunter, A. 1991. Traits that distinguish outbreaking and nonoutbreaking macrolepidoptera feeding on northern hardwood trees. Oikos 60:275–282.

Hunter, A. F., and J. S. Elkinton. 2000. Effects of synchrony with host plant on populations of a spring-feeding lepidopteran. Ecology 81:1248–1261.

Hunter-Fujita, F. R., P. F. Entwistle, and H. R. Evans. 1998. Insect Viruses and Pest Management. John Wiley and Sons.

Johnson, D. M., A. M. Liebhold, O. N. Bjornstad, and M. L. McManus. 2005. Circumpolar variation in periodicity and synchrony among gypsy moth populations. J. Anim. Ecol. 74:882–892.

Jones, C. G., R. S. Ostfeld, M. P. Richard, E. M. Schauber, and J. O. Wolff. 1998. Chain reactions linking acorns to gypsy moth outbreaks and Lyme disease risk. Science 279:1023–1026.

Keeling, M. J., and P. Rohani. 2008. Modeling infectious diseases in humans and animals. Princeton University Press.

Kendall, B. E., C. J. Briggs, W. W. Murdoch, P. Turchin, S. P. Ellner, et al. 1999. Why do populations cycle? A synthesis of statistical and mechanistic modeling approaches. Ecology 80:1789–1805.

Kraaijeveld, A., J. Ferrari, and H. Godfray. 2002. Costs of resistance in insect-parasite and insect-parasitoid interactions. Parasitology 125:S71–S82. Meeting Parasite Variation: Immunological and Ecological Significance, London, England, Sept 14, 2001.

Liebhold, A., and N. Kamata. 2000. Are population cycles and spatial synchrony a universal characteristic of forest insect populations? Population Ecology 42:205–209.

Liebhold, A. M., R. Plymale, J. S. Elkinton, and A. E. Hajek. 2013. Emergent fungal entomopathogen does not alter density dependence in a viral competitor. Ecology 94:1217–1222.

McCullagh, P., and J. A. Nelder. 1989. Generalized linear models, vol. 2. Chapman and Hall London.

McManus, M., and C. C. Doane. 1981. The gypsy moth: Research toward integrated pest management. Technical Bulletins 158053, United States Department of Agriculture, Economic Research Service.

Moreau, G., and C. J. Lucarotti. 2007. A brief review of the past use of baculoviruses for the management of eruptive forest defoliators and recent developments on a sawfly virus in Canada. For. Chron. 83:105–112.

Moreau, G., C. J. Lucarotti, E. G. Kettela, G. S. Thurston, S. Holmes, C. Weaver, D. B. Levin, and B. Morin. 2005. Aerial application of nucleopolyhedrovirus induces decline in increasing and peaking populations of Neodiprion abietis. Biol. Control 33:65–73.

Murray, K. D., and J. S. Elkinton. 1989. Environmental contamination of egg masses as a major component of transgenerational transmission of gypsy-moth nuclear polyhedrosis virus (LdMNPV). J. Invertebr. Pathol. 53:324–334.

Myers, J. H. 1988. Can a general hypothesis explain population cycles of forest Lepidoptera? Adv. Ecol. Res. 18:179–242.

Myers, J. H. 1993. Population outbreaks in forest Lepidoptera. Amer. Sci. 81:240–251.

Narciso, D. 2014. Some Central Ohioans Object to Gypsy-moth Spraying. The Columbus Dispatch, 17 November.

Nealis, V. G. 1991. Parasitism in sustained and collapsing populations of the jack pine budworm, Choristoneura pinus pinus Free. (Lepidoptera: Torticidae), in Ontario, 1985-1987. Can. Entomol. 123:1065.

Nolan, I. 2015. State agency plans second meeting to update gypsy moth plan. Island Free Press: Hatteras and Ocracoke Island News, 27 October 2015.

Páez, D. J., A. Fleming-Davies, and G. Dwyer. 2015. Effects of pathogen exposure on life history variation in the gypsy moth (Lymantria dispar). J. Evol. Biol. 28:1828–1839.

Parker, B. J., B. D. Elderd, and G. Dwyer. 2010. Host behaviour and exposure risk in an insect-pathogen interaction. J. Anim. Ecol. 79:863–870.

Plummer, M., N. Best, K. Cowles, and K. Vines. 2006. Coda : Convergence diagnosis and output analysis for MCMC. R News 6:7–11.

Podgwaite, J. D., N. R. Dubois, R. C. Reardon, and J. Witcosky. 1993. Retarding outbreak of low-density gypsy-moth (Lepidoptera: Lymantriidae) populations with aerial applications of gypchek and Bacillus thuriengensis. J Econ Entomol 86:730–734.

Podgwaite, J. D., R. C. Reardon, G. S. Walton, and J. Witcosky. 1992. Efficacy of aerially-applied Gypchek against gypsy-moth (Lepidopera: Lymantriidae) in the Appalachian Highlands. J. Entomol. Sci. 27:337–344.

Reilly, J. R., and B. D. Elderd. 2014. Effects of biological control on long-term population dynamics: identifying unexpected outcomes. J. Appl. Ecol. 51:90–101.

Rossiter, M. C. 1991. Environmentally-based maternal effects – a hidden force in insect population-dynamics. Oecologia 87:288–294.

Sasaki, A., and H. C. J. Godfray. 1999. A model for the coevolution of resistance and virulence in coupled host-parasitoid interactions. Proc. R. Soc. B 266:455–463.

Schreiber, S. J., R. Burger, and D. I. Bolnick. 2011. The community effects of phenotypic and genetic variation within a predator population. Ecology 92:1582–1593.

Skaller, P. M. 1985. Patterns in the distribution of gypsy moth *(Lymantria dispar)* egg masses over an 11 year population cycle. Environ. Entomol. 14:106–117.

Turchin, P. 2003. Complex Population Dynamics: A Theoretical/empirical Synthesis. Monographs in population biology. Princeton University Press.

Walter, J. A., D. M. Johnson, P. C. Tobin, and K. J. Haynes. 2015. Population cycles produce periodic range boundary pulses. Ecography 38:1200–1211.

Watanabe, H. 1987. The host population. Pages 71–112 in J. Fuxa and Y. Tanada, eds. Epizootiology of insect diseases. Wiley: New York.

Williams, D. W., R. W. Fuester, W. W. Metterhouse, R. J. Balaam, R. H. Bullock, R. J. Chianese, and R. C. Reardon. 1990. Density, size and mortality of egg masses in New Jersey populations of the gypsy moth (Lepidoptera, Lymantriidae). Environ. Entomol. 19:943–948.

Woods, S. A., and J. S. Elkinton. 1987. Bimodal patterns of mortality from nuclear polyhedrosis-virus in gypsy-moth (Lymantria dispar) populations. J. Invertebr. Pathol. 50:151–157.

Yoshida, T., S. P. Ellner, L. E. Jones, B. J. M. Bohannan, R. E. Lenski, and N. G. Hairston, Jr. 2007. Cryptic population dynamics: Rapid evolution masks trophic interactions. PLoS Biology 5:1868–1879.

Yoshida, T., L. E. Jones, S. P. Ellner, G. F. Fussmann, and N. G. Hairston. 2003. Rapid evolution drives ecological dynamics in a predator-prey system. Nature 424:303–306.

Zbinden, M., C. R. Haag, and D. Ebert. 2008. Experimental evolution of field populations of Daphnia magna in response to parasite treatment. J. Evol. Biol. 21:1068–1078.

